# Deciphering the diversity and sequence of extracellular matrix and cellular spatial patterns in lung adenocarcinoma using topological data analysis

**DOI:** 10.1101/2024.01.05.574362

**Authors:** Iris H.R. Yoon, Robert Jenkins, Emma Colliver, Hanyun Zhang, David Novo, David Moore, Zoe Ramsden, Antonio Rullan, Xiao Fu, Yinyin Yuan, Heather A. Harrington, Charles Swanton, Helen M. Byrne, Erik Sahai

**Author notes:** these authors contributed equally (Robert Jenkins, Emma Colliver, Hanyun Zhang).

## Abstract

Extracellular matrix (ECM) organization influences cancer development and progression. It modulates the invasion of cancer cells and can hinder the access of immune cells to cancer cells. Effective quantification of ECM architecture and its relationship to the position of different cell types is, therefore, important when investigating the role of ECM in cancer development. Using topological data analysis (TDA), particularly persistent homology and Dowker persistent homology, we develop a novel analysis pipeline for quantifying ECM architecture, spatial patterns of cell positions, and the spatial relationships between distinct constituents of the tumour microenvironment. We apply the pipeline to 44 surgical specimens of lung adenocarcinoma from the lung TRACERx study stained with picrosirius red and haematoxylin. We show that persistent homology effectively encodes the architectural features of the tumour microenvironment. Inference using pseudo-time analysis and spatial mapping to centimetre scale tissues suggests a gradual and progressive route of change in ECM architecture, with two different end states. Dowker persistent homology enables the analysis of spatial relationship between any pair of constituents of the tumour microenvironment, such as ECM, cancer cells, and leukocytes. We use Dowker persistent homology to quantify the spatial segregation of cancer and immune cells over different length scales. A combined analysis of both topological and non-topological features of the tumour microenvironment indicates that progressive changes in the ECM are linked to increased immune exclusion and reduced oxidative metabolism.

## Introduction

Cancer is a major cause of mortality globally, with lung cancer accounting for 1.8M deaths annually [1]. The lethal progression of lung tumours from benign to metastatic is linked to both tumour cell-intrinsic changes and paracrine interactions between cancer cells and the tumour microenvironment (TME). Within the TME, cancer cells interact with various cell types, including leukocytes, endothelial cells, cancer-associated fibroblasts (CAFs), and the extracellular matrix (ECM). The ECM directly influences tumour cell growth, invasion, and resistance to chemotherapy [2–4]. The ECM is also proposed to act as a barrier for immune cell access to tumours, thus modulating anti-tumour immunity and blunting responses to immunotherapy [4]. Developing a clear understanding of the roles of the ECM and the relative spatial positioning of cells in tumour progression and therapy responses requires quantitative descriptors that can be linked to other classes of data, such as transcriptomes, mutational profiles, and patient outcomes. Although certain metrics of ECM organisation, such as density and alignment, have been developed and have been shown to have prognostic value in many cancer types [5–10], there is no simple method for combining quantification of ECM and cellular information.

Recent years have seen the development of a number of tools to analyse the spatial relationship between cells, including neighbourhood analysis [11, 12] and topological tumour graphs [13]. Although versatile and powerful, neighbourhood analysis is intended for thecharacterisation of cell clusters, not ECM fibres [11, 12]. Moreover, it cannot readily incorporate information about how patterns vary as a function of length scale. Topological data analysis (TDA), a field of computational mathematics, provides a suite of methods for studying spatial patterns. A popular technique of TDA, called persistent homology (PH), takes as input the point cloud of cell locations extracted from image data. It then builds a nested sequence of simplicial complexes, which are higher-dimensional analogues of graphs (Fig. 1B) [14–16]. The sequence of simplicial complexes is indexed by a scale parameter (denoted ε in Fig. 1B). PH then examines the birth and death of spatial features, such as clusters, loops, and voids across the simplicial complexes. The output of PH, called persistence diagrams (PD), summarises the parameters at which the spatial features appear and die (see *Methods*- dimension 0 PDs summarize the parameters at which components merge, analogous clustering, while dimension 1 PDs summarize the formation and filling-in of loops (Fig. 1B)). Persistence diagrams thus provide a multiscale and multidimensional descriptor of spatial patterns. The persistence diagrams can be vectorised for machine learning tasks [14–16]. The utility of PH is evident in its application to cellular patterns arising in histology images of breast and prostate cancer [17–23]. Moreover, the ability to describe loops and higher-dimensional features across length scales makes TDA well-suited to ECM analysis.

**Figure 1.**
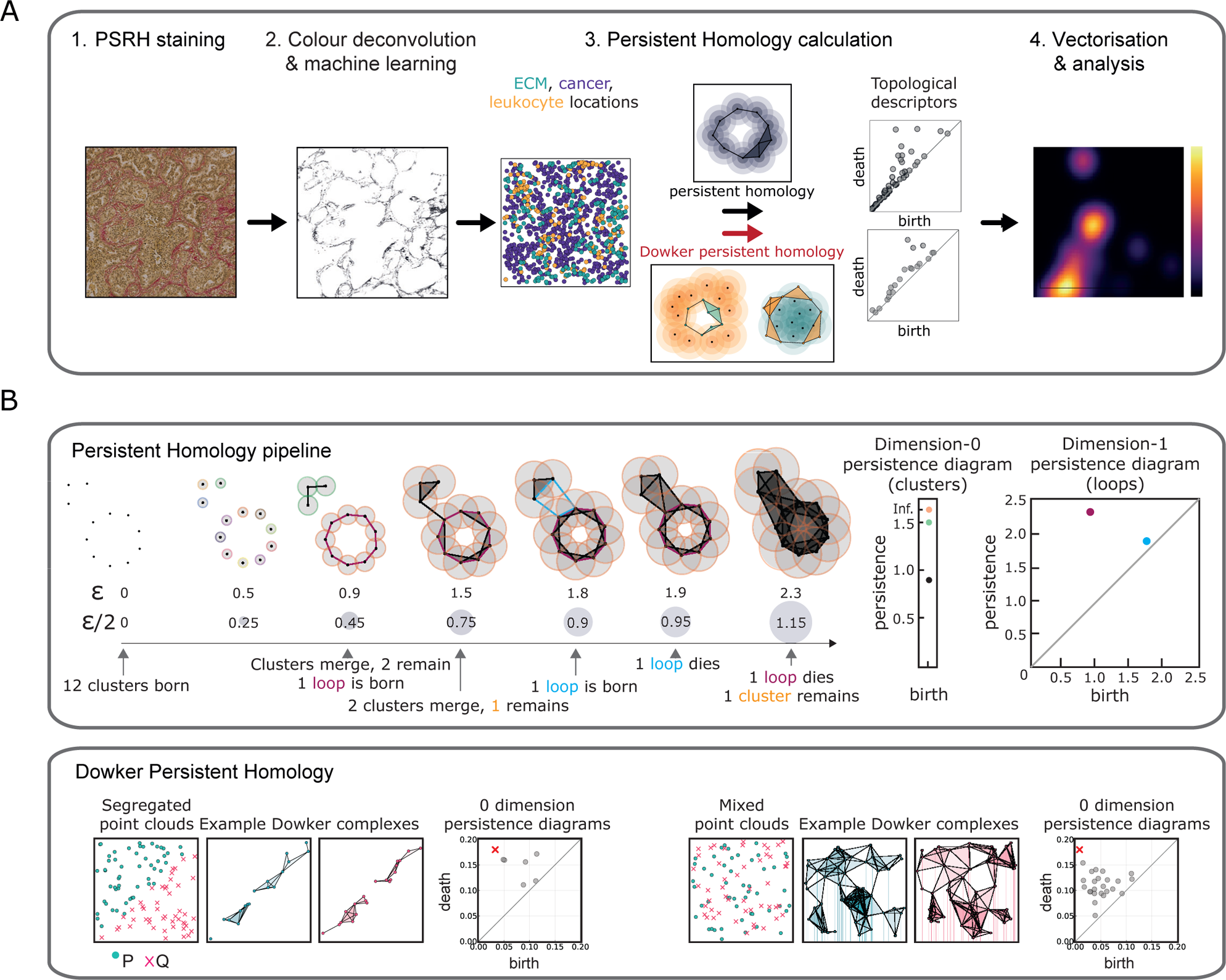
Pipeline for constructing topological feature vectors. **A.** Given a PSRH image, we obtain an ECM image via colour deconvolution and sample points from the ECM image according to pixel intensity. We extract locations of cancer cells and leukocytes via an in-house deep neural network. We use tools in topological data analysis, namely persistent homology and Dowker persistent homology, which result in topological descriptors, called persistence diagrams, that summarize the presence of structural elements (connected components and loops). The topological descriptors are vectorised and analysed via dimensionality reduction. **B.** Upper panel: Persistent homology pipeline. Given a point cloud and a parameter *ε*, one creates a graph by placing a line between two points if their distance is at most *ε*. This process is equivalent to placing balls of radius 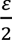 around each point and creating a line between two points if the balls intersect. One then fills in any triangles, tetrahedra, and higher-dimensional objects of the graph. We refer to the resulting object as a simplicial complex at parameter *ε*. Persistent homology records the birth and death of topological features as the parameter *ε* increases. The 0-dimensional persistence diagram records the parameters at which clusters merge. The 1-dimensional persistence diagram summarize the parameters at which loops emerge and die. Lower Panel: Example output of Dowker persistent homology on a pair of point clouds. The 0-dimensional Dowker persistent homology summarizes boundaries and shared connected components between a given pair. The 0-dimensional Dowker persistence diagram from segregated point clouds lack points near the origin in comparison to the 0-dimensional Dowker persistence diagram from mixed point clouds.

While PH provides automatic and effective methods for summarising spatial patterns, its application is limited to the analysis of a single entity within a system – e.g., either ECM, cancer cells, or leukocytes. In particular, it does not elucidate how features associated with two systems or populations are related. Given two systems, Dowker persistent homology (Dowker PH) [24, 25] studies spatial patterns by forming simplicial complexes on one system whose construction depends on proximity to the other system (see *Methods*). While its closely related cousin Witness complexes enjoyed popularity as a tool to quickly and approximately compute persistent homology, only recently have Dowker complexes been utilised to study relational data, such as protein-ligand binding, PDF files and parser messages [26–28]. The outputs of Dowker PH, such as Dowker persistence diagrams (see *Methods*), which summarise the length scale at which shared clusters or loops form and extinguish, can be made compatible with conventional statistics and machine learning via vectorisations [29].

Understanding how tumours develop and evolve over time is crucial for improving cancer treatment. TRACERx (TRAcking non-small cell lung Cancer (NSCLC) Evolution through therapy (Rx); ClinicalTrials.gov: NCT01888601) is a prospective study of tumour evolution in patients with early-stage resectable NSCLC. The TRACERx study is tracking the progression of tumours in a scheduled full cohort of 842 patients, including multi-region tissue sampling and detailed genomic, transcriptomic, and tumour microenvironment characterisation [30–36]. In this work, we study spatial patterns, as determined by persistent homology and Dowker persistent homology, in a subset of lung adenocarcinoma from the TRACERx study lung tumours (see *Methods*). In particular, we use persistent homology to study the architectural features of ECM, cancer cells, and leukocytes. Our analysis shows that TDA provides robust features that capture the patterns and organisation of ECM fibres, cancer cells, and leukocytes. Moreover, we deploy Dowker persistent homology in a novel way to quantify colocalization and exclusion between two constituents, with particular utility for classifying patterns of spatial segregation between cancer cells and leukocytes, termed immune exclusion. The application of ‘pseudo-time’ analysis suggests likely routes of transition between patterns of tissue organisation. Moreover, we demonstrate that spatial patterns inferred to be closely linked in time are also physically proximal in tissue, arguing for graded and progressive changes in tissue architecture. Lastly, we integrate spatial pattern information with transcriptomic data to derive a wholistic view of how the tumour microenvironment changes as lung adenocarcinomas evolve.

## Results

### Overview of pipeline for extracting topological features of lung adenocarcinoma

To extract quantitative features of ECM, cancer cell, and leukocyte organisation in lung cancer, we established a data acquisition and analysis pipeline (Fig. 1). Diagnostic lung adenocarcinoma slides were stained with PicroSirius Red (PSR), highlighting fibrillar collagen I and III, and haematoxylin, which stains cell nuclei (we refer to this combined stain as PSRH). Following scanning, a colour deconvolution step was performed to separate the PSR ECM stain. In addition, a fully convolutional network Symmetric Distance Regularized Dense Inception Neural network (S-DRDIN) was used to identify cancer cells and leukocytes from the PSRH images (precision 0.90 and 0.84, respectively). The algorithm was also trained to identify red blood cells and anthracotic particulate pollution with high accuracy (0.92 and 0.94, respectively). The precision of identifying fibroblasts and necrotic cells was below 0.7 and these features were not used for subsequent topological data analysis. For ease of data processing, the large whole slide images (typically 15mm x 30mm) were divided into regions of interest (ROIs) of 4000 pixels x 4000 pixels, corresponding to 878µm x 878μm. Point clouds representing the cancer cell and leukocyte distributions were extracted from the machine learning method mentioned above, and point clouds representing the ECM were obtained by sampling from the ROIs (SI Fig. 1A-C). Persistent homology was used to generate two topological descriptors, the 0-dimensional and 1-dimensional persistence diagrams. The 0-dimensional persistence diagram summarises the merging of clusters in the data (Fig. 1B), while the 1-dimensional persistence diagram summarises the birth and death of loops (Fig. 1B). These topological descriptors were derived for the ECM, cancer cells, and leukocytes separately. The 0-dimensional and 1-dimensional persistence diagrams were then vectorised to yield 20-dimensional and 400-dimensional vectors, respectively (see *Methods* and Supp. Fig. 1D). We refer to the resulting vectors as 0-dimensional and 1-dimensional *PH features*. Dowker persistent homology was applied to pairs of ECM points, cancer cells, and leukocytes. Similar to persistent homology, Dowker persistent homology also results in two topological descriptors; the 0-dimensional Dowker persistence diagram summarises the birth and death of neighbouring clusters between the pair (Fig 1C, top) and the 1-dimensional Dowker persistence diagram summarises the birth and death of shared loops (Fig 1C, bottom). The Dowker 0-dimensional and 1-dimensional PDs were vectorised yielding 400-dimensional vectors. Together, these analyses summarise both the individual cluster and loop features of ECM, cancer cell, and leukocyte distributions, and their spatial inter-relationships.

### Persistent homology features are robust to image acquisition method

An overarching goal in developing powerful data analysis methodology should be its independence from noise and the data acquisition technology. The success of PH across the sciences [37–39] is, in part, due to the stability theorem [40, 41] which states that topological features are robust to small noise perturbations. We tested the robustness of TDA features with respect to changes in input image resolution on 100 randomly selected ROIs. For each ROI, we first created lower-resolution ECM images by progressively downsizing the image and then resizing it to have the same number of pixels as the original image. We sampled point clouds representing ECM from the images of varying resolutions and computed the difference between the 1-dimensional persistence images from the lower-resolution ECM images and the persistence images from the original ECM images (via the Frobenius norm of the two persistence images). As shown in Supp. Fig. 2A, the average difference between two samplings of the same ROI was approximately 1.6 × 10^−5^ regardless of the extent of downsizing (upto 16-fold downsizing was tested). For reference, the difference between persistence images in two distinct ROIs is 7.9 × 10^−5^.

These analyses show that the outputs of persistent homology will be robust when applied to images with pixel sizes in the range 0.2-2μm, which comfortably spans the resolution of light microscopy-based methods. To explore this directly, we stained serial lung sections with PSRH and an antibody against collagen I. The antibody staining was acquired using the AKOYA Phenocycler platform with a resolution of 0.4μm/pixel, which contrasts with 0.22μm/pixel for the PSRH images. Supp. Fig. 2B shows the similarity of the persistence images from ECM stains obtained from two different data acquisition methods.

### Persistent homology features reliably discriminate patterns of ECM organisation

Having established that TDA could be applied to ECM, we tested if it could effectively distinguish different ECM patterns in lung cancer. Four hundred 878 x 878μm^2^ regions of interest (ROIs) were selected based on diversity of ECM and cellular organisation from 44 different LUAD diagnostic samples from the lung TRACERx study [30, 31] (see *Methods*). Following colour deconvolution to separate the collagen, we sampled point clouds according to the density of ECM (Supp. Fig. 1A). We computed the 0-dimensional and 1-dimensional persistence diagrams which respectively summarize the clusters and loops in ECM. Their vectorizations, referred to as 0-dimensional and 1-dimensional PH features, were concatenated to a 420-dimensional vector (Fig. 2A).

**Figure 2.**
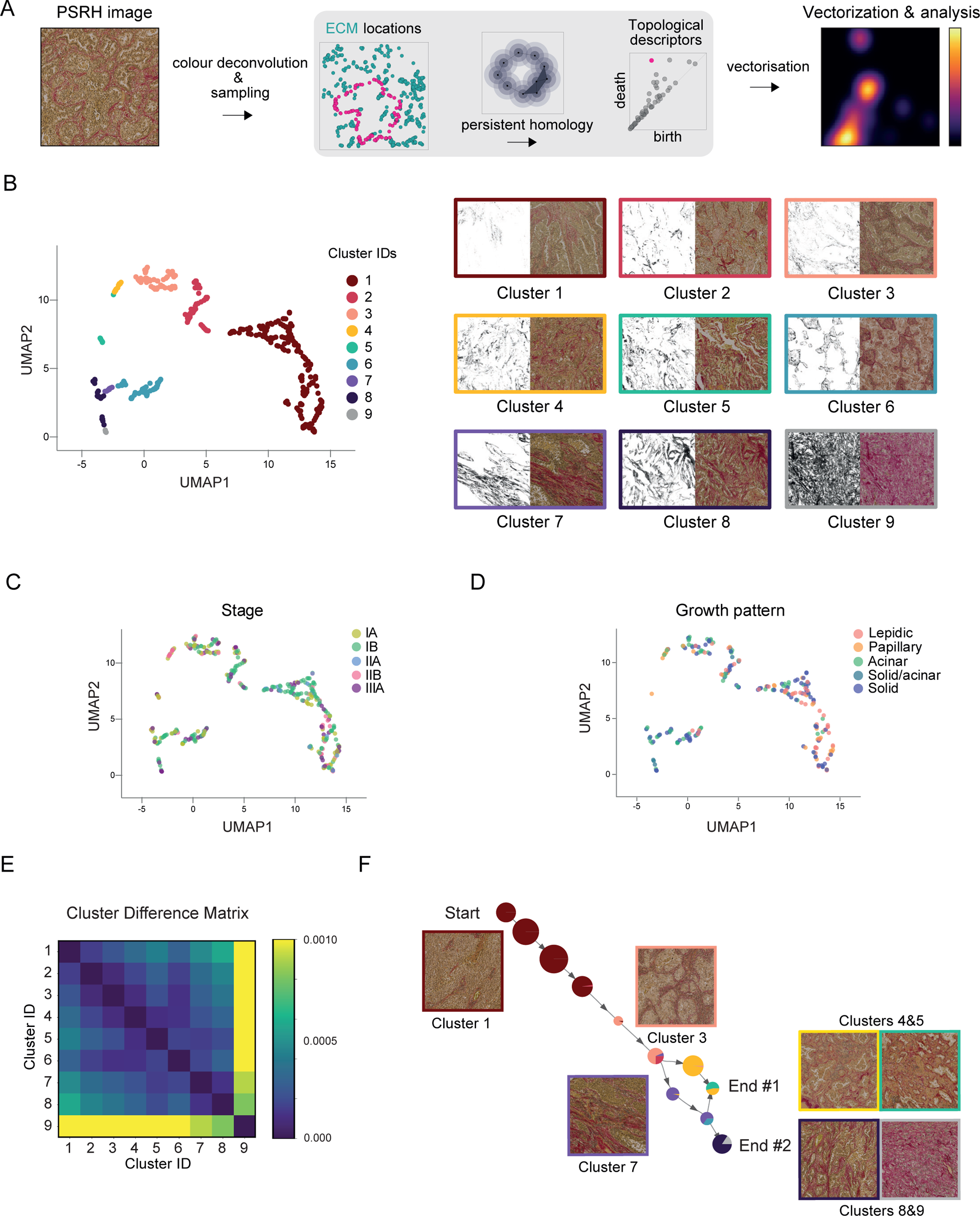
Persistent homology features reveal patterns of ECM architecture. **A.** Construction of PH feature vectors. From each PSRH image, we only consider the ECM points. We apply persistent homology to the ECM points and obtain 0-dimensional persistence diagrams and 1-dimensional persistence diagrams, which summarize the birth and death of clusters and loops. The location of the loop highlighted in magenta in the persistence diagram is indicated by the magenta dot. We vectorised the persistence diagrams and concatenated the resulting vectors. All analysis was performed on the concatenated vector, which we refer to as ECM PH feature. **B.** (Left) Applying UMAP on the ECM PH feature reveals 8 clusters, labelled 1 to 8. Points not assigned to clusters were manually identified to occupy a similar area of parameter space and designated cluster 9. (Right) Representative images of clusters. Each image is 878µm x 878µm. **C.** Tumour stage is overlaid on UMAP of ECM PH features. **D.** Histological growth pattern is overlaid on UMAP of ECM PH features. **E.** Distance matrix shows pairwise distances between the clusters computed by the Euclidean distance between the average PH feature vectors of each cluster. **F**. Pseudo-time analysis demonstrates likely trajectories from Cluster 1 to terminal endpoints at clusters 4&5 and 8&9. Circles represent clusters predicted by the pseudo-time PARC algorithm, with proportions of clusters from Figure 2B shown.

To analyse the PH feature vectors, we performed both UMAP and PCA dimensionality reduction. Applying UMAP on the ECM PH feature revealed 8 clusters, labelled 1 to 8. Points that were not assigned to the eight clusters occupied a similar region in the parameter space and were designated cluster 9. Fig 2B shows exemplar PSRH and ECM stains for each cluster and provides visual confirmation of the ability of our pipeline to group similar ECM patterns together. Visual inspection revealed that the first dimension of the UMAP corresponds primarily to ECM abundances in the ROIs, with cluster 1 having sparse ECM and clusters 7-9 having dense ECM. The relative ability of 0-dimensional and 1-dimensional PH feature vectors to discriminate clusters is shown in Supp. Fig. 3B. PCA does not generate the discrete clusters visible in the UMAPs. However, it has the advantage that the principal components can be visualised as persistence images. The weighting of different regions of the 1-dimensional persistence images is shown in Supp. Fig. 3C. The first principal component (PC1) weighting penalises small loops that fall within the dark patch (marked with white asterisk), while PC2 favours larger loops that fall within the yellow/white patch (marked with orange asterisk). We associate low PC1 coordinates with ECM abundance as this generates many small loops (Supp. Fig. 3D). High second principal component (PC2) coordinates are indicative of slightly larger loops, with low PC2 values reflecting a lack of such loops, which can arise from ECM that is uniformly distributed or from ECM whose structural organization occurs at a larger Supp. Fig. 3D). The interpretability of persistence images, coupled with the interpretability of the eigenvectors of PCA, enable the identification of the dominant ECM features in different ROIs. Analysis of intra-tumour heterogeneity of ECM illustrated that the majority of tumours contained ECM patterns that fell into different clusters (Supp. Fig. 3E). These analyses demonstrate the ability of persistent homology to describe ECM features, with downstream UMAP analysis identifying recurring classes of ECM organisation and PCA providing interpretable, low-dimensional description of the tissue pattern.

To understand more precisely how the PH features relate to traditional ECM features, we subjected the same 400 ROIs to analysis of discrete fibres using CT-FIRE, gap analysis, and texture feature algorithms (Fu, Jenkins *et al.*, in preparation). UMAP on these non-topological features shows less defined clustering than that based on PH features (Supp. Fig. 3F). Overlaying the cluster identities derived from the PH analysis (Fig. 2B) on the UMAP of non-topological features showed similar relationships among the 400 ROIs, albeit less well-defined (Supp. Fig. 3F). Supp. Fig. 3G shows how individual features of ECM density, gaps, fibre persistence, and fractal dimension relate to the UMAP of PH features (Fig. 2B). These suggest that cluster 1 has low ECM density, large circular gaps, and low fractal dimension, and that cluster 8 has opposite features. Together, these analyses demonstrate that PH features capture multiple aspects of the spatial patterns of ECM and that clustering on PH features leads to more well-defined clusters in comparison to non-topological features.

### Linkage of ECM patterns to clinical parameters

Next, we sought to understand the relationship between the PH features of ECM, a pathologist’s classification of lung cancer architecture, and patient characteristics including age and gender. There was no clear association between ECM organisation and stage (Fig. 2C). There were also no clear links between ECM PH features and age, gender, smoking status, tumour mutational burden, or *KRAS* or *TP53* mutation (Supp. Fig. 4A&B). Overlaying histological growth pattern information revealed that lepidic and papillary histologies were more common in cluster 1 (Fig. 2D). Also, the same histological growth pattern could have distinct ECM organisation, with solid histology spanning cluster 1 to 8. Thus, PH features of ECM can capture information that is additional to histological growth patterns.

Pollution has been linked to the development and progression of lung cancer [33]. Of note, particulate pollution, such as PM2.5, can be retained in the lungs for long periods of time and is visible as anthracotic pigment in histological sections. Therefore, we explored the association between ECM organisation and regions containing particulate pollutants. The analysis revealed that clusters 7 and 9 had elevated levels of particulate matter, suggesting that it may promote the transition of lung cancer tissue to fibrotic states (Supp. Fig. 4C).

### Inference of transitions between patterns of ECM organisation

The analysis described above identifies quantitatively distinct classes of ECM organisation. We investigated the relative distance of the nine clusters by computing the Euclidean distance between the average feature vectors of each cluster. The distance matrix reveals similarities between many of the clusters, with a dominant diagonal trend evident: cluster 1 is most similar to cluster 2, cluster 2 being most similar to clusters 1 & 3, and so on (Fig. 2E). Consistent with the HDBSCAN analysis, cluster 9 is an outlier, although it shares some similarity with cluster 8. The relative similarity of the feature vectors of the clusters prompted us to explore the likely route of transition between different patterns of ECM organisation. Pseudotime analysis revealed that the most likely sequence of events could be described with 11 stages, initiating with cluster 1 and having two possible end states (Fig. 2F). The cluster 4/5 end state could be reached either via progression from cluster 3 or cluster 6/7. The cluster 8/9 end state was reached via sequential transitions from cluster 1 to 2/3, 2/3 to 7, and then to clusters 8/9. These analyses suggest that ECM changes in the lung begin along a common trajectory, with some bifurcations leading to a fully fibrotic state with high ECM density and few cancer cells.

### Persistent homology analysis of large tissue areas suggests ECM changes in a progressive manner

To understand the spatial relationship and proximity among tissues with different ECM organisation, we selected 20 diagnostic LUAD sections (∼15mm x 30mm) and decomposed them into 9,382 ROIs of size 878μm x 878μm. For each ROI, we computed the 0-dimensional and 1-dimensional PH features and concatenated them into a combined PH feature vector. Each ROI was then assigned to a cluster from Fig. 2B based on the shortest Euclidean distance between its PH vector and the average feature vector of the clusters in Fig. 2B. Fig. 3A shows an example of a whole slide PSR image overlaid with colour-coding to reflect the assigned cluster (further examples are shown in Supp. Fig. 5). Similar to Fig. 2, UMAP analysis suggested gradual transitions between ECM patterns (Fig. 3B). The use of whole slide images also enabled the analysis of tumour-adjacent, non-cancerous tissue via PH features. Fig. 3C shows that the majority of non-cancerous ROIs fell within cluster 1 (Fig. 3B), which is consistent with the pseudo-time analysis being initiated from cluster 1 (Fig. 2F). However, there were many non-cancerous regions that mapped to other clusters. Visual inspection of the ROIs revealed that even though the regions lacked cancer cells, the lung architecture was not entirely normal in these regions, with some clearly exhibiting increased PSR signal (Fig. 3C). These data indicate that transitions in ECM organisation can be observed in tissue adjacent to the tumour.

**Figure 3.**
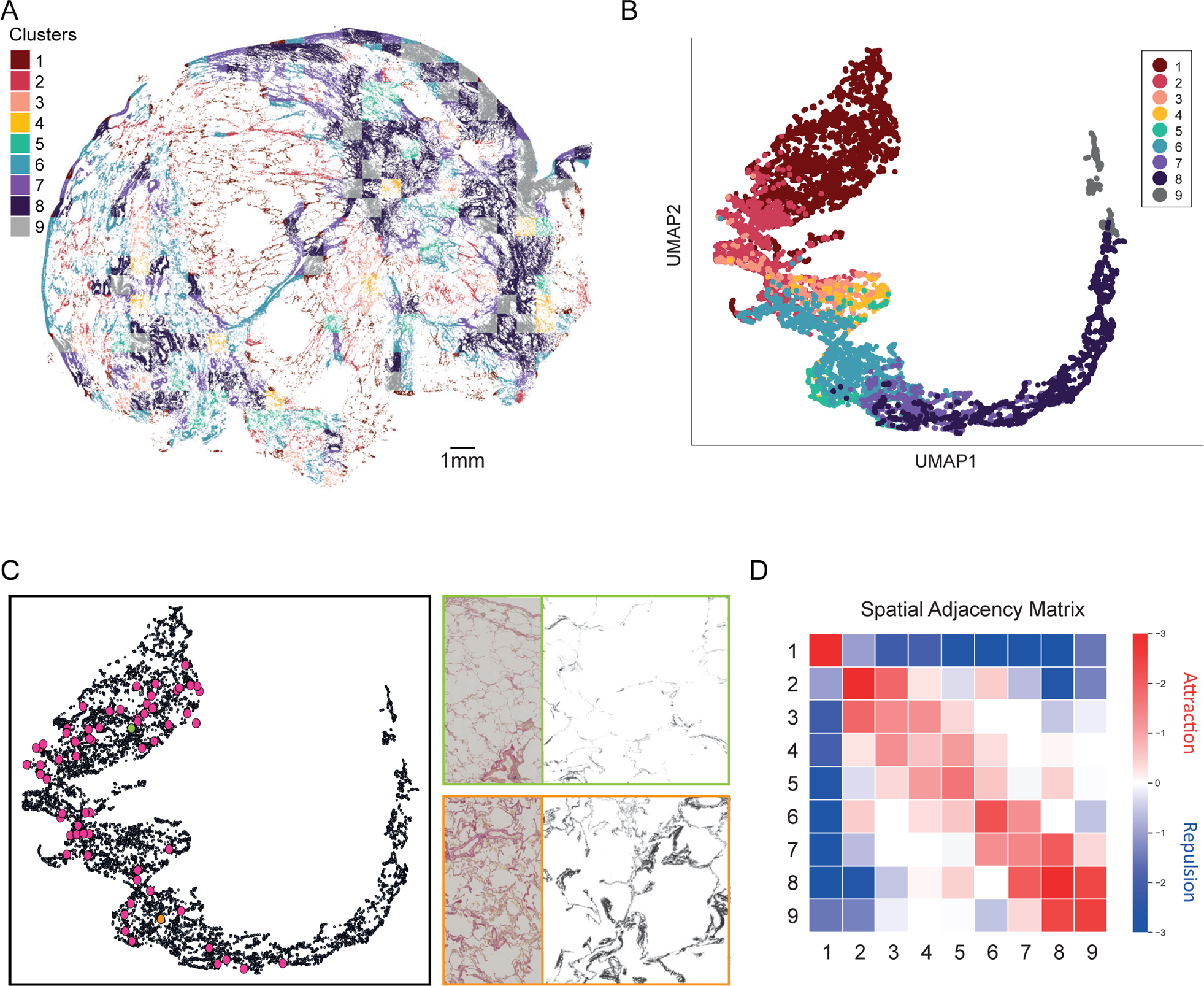
Persistent homology analysis of large tissue areas suggests ECM changes in a progressive manner. **A.** Visualization of a whole-slide tumour, with smaller regions coloured by the closest cluster in Fig. 2B. The whole-slide ECM stain was split into tiles of size 878μm x 878μm. For each tile, we computed the Euclidean distance between its PH feature vector and the average feature vector of each cluster in Fig. 2B. We then assigned the cluster with the shortest distance and coloured the tile according to the assigned cluster. **B.** UMAP of 9382 regions analysed from 20 whole-slide images. Colours indicate the assignment to the previously described clusters (Fig. 2B). **C.** Non-cancerous adjacent regions are highlighted in the UMAP plot, together with two exemplar images of adjacent tissue. Green dot and box highlight non-cancerous tissue falling cluster 1 and orange dot and box highlight non-cancerous tissue grouping in cluster 6. Coloured PSRH image is 2.5mm x 5mm and grayscale image showing only collagen is 878µm x 878µm. **D.** Matrix shows the relative ‘adjacency’ of different clusters to each other. For each pair of clusters, the frequency of neighbouring ROIs was determined and ranked against 1999 randomly generated spatial distributions of ROIs to provide a p-value. A red colour indicates adjacency with a greater than expected frequency, a blue colour indicates a lower expected frequency with values given on a log_10_ scale.

We then examined whether certain patterns of ECM are typically located adjacent to other patterns of ECM. Given two clusters, we computed their neighbouring frequency, defined as the number of pairs of ROIs from each cluster that shared an edge. This frequency was compared to the neighbouring frequency after randomly shuffling the assigned cluster of each ROI (see *Methods*). The result was a set of pairwise comparisons indicating whether two clusters were neighbours with a greater than expected, expected, or less than expected frequency, as determined by calculating p-values from the proportion of randomisations above or below the observed data (Fig. 3D). The significance of the p-value is indicated by darkness of colour. Greater than expected interactions are shown in red whilst lower than expected interactions are shown in blue. The diagonal pattern (from top left to bottom right) of greater than expected interaction indicates that an ECM region with a given cluster identity is likely to neighbour another region of the same identity, or with a cluster identity one higher or lower than its own. For example, a cluster 8 region is likely to have neighbours from clusters 7, 8, or 9. An exception is cluster 1, that is only likely to have neighbours in cluster 1. This means that there are typically large areas of cluster 1. However, once the ECM begins to transition towards increased ECM density, then regions with similar pattern are likely to be located adjacent to each other.

### Combined PH features capture patterns of cancer cells and leukocyte distribution

Immune cells play critical roles in cancer biology. Therefore, we focussed next on describing the distribution of cancer cells and leukocytes in the tumour regions described in Figure 2. computed four PH features: 0-dimensional and 1-dimensional PH features from cancer cells, and 0-dimensional and 1-dimensional PH features from leukocytes (identified via the application of machine learning to PSRH images – see *Methods*). We concatenated the four feature vectors to one large combined PH feature vector and performed UMAP (Fig. 4A). Visual inspection confirmed that the clusters capture different patterns of tissue organisation. UMAP on each of the four PH features is shown in Supp. Fig. 6A, with colouring indicating the clusters assigned from the analysis of the combined PH feature vector in Fig. 4A. The clustering derived from the combined PH feature vector is most clearly retained in the cancer cell UMAPs, suggesting that cancer cell distribution plays a dominant role in the analysis of the combined PH feature vectors. Of note, clusters 1, 2, and 10 identified regions within which cancer cells and leukocytes were excluded from one another. These clusters are also characterised by higher leukocyte densities, fewer cancer cells, and distinct ECM organisation (Fig. 4B and Supp. Fig. 6B). Cross-referencing with histological growth patterns revealed that acinar histology was predominantly found in clusters 1 and 2, and lepidic and papillary histologies were dominant in clusters 4, 5, 7, and 8 (Supp. Fig. 6B). No clear linkage was observed with KRAS or TP53 mutations (Supp. Fig. 6B). Principal component analysis on the combined PH feature indicates that the principal components encode relative abundance (PC1) and segregation (PC2) of cancer cells and leukocytes (Supp. Fig. 6C). Together, these analyses demonstrate the ability of PH features to reliably identify regions with similar spatial patterns of cancer cells and leukocytes, and they further suggest an association between ECM and cellular patterns.

**Figure 4.**
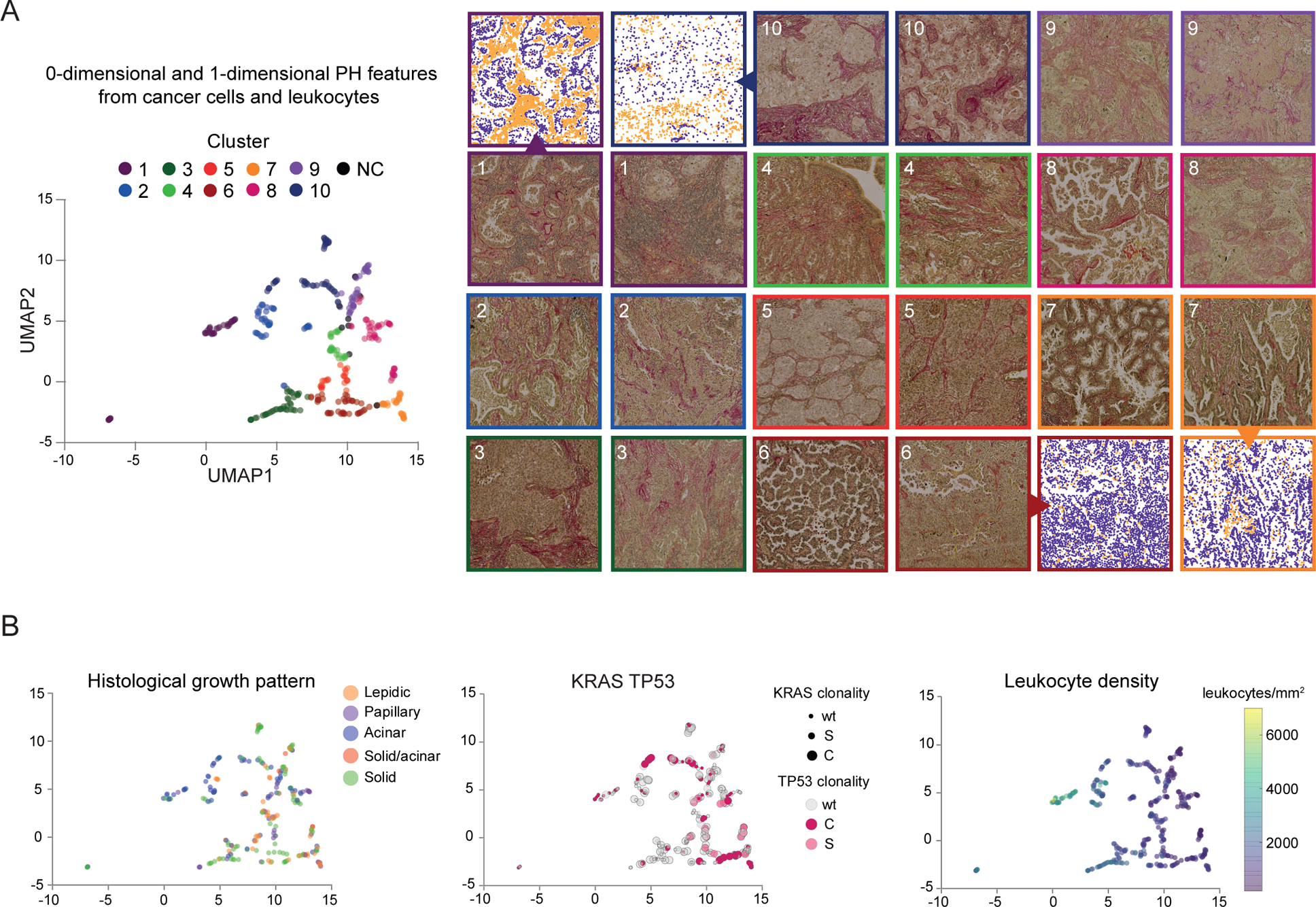
Combined PH features of cancer cells and leukocytes capture patterns of cancer cell and leukocyte distribution. **A.** Four PH features (dimension-0 and dimension-1 PH features on cancer cells and leukocytes) are concatenated to form a combined PH feature vector. (Left) Applying UMAP on the combined PH features reveals 10 clusters, labelled 1 to 10 and assigned distinct colours. Points not assigned to clusters were distributed in different areas of parameter space and are labelled NC. (Right) Representative examples from each of the clusters. The numbering and border colour indicate the cluster to which the image belongs. Each image is 878µm x 878µm. **B.** UMAP of combined PH feature overlaid with (left) histological growth pattern, reflected in dot colours, (middle) *KRAS* mutation (dot size, wt – wildtype, C – clonal, S – sub-clonal) and *TP53* mutation (dot colour, wt – wildtype, C – clonal, S – sub-clonal), and (right) machine learning-derived leukocyte densities.

### Dowker PH features enable quantification of immune cell exclusion from cancer cells

By concatenating the PH features from cancer cells and leukocytes, we were able to identify characteristic patterns of immune cells in tumours. However, concatenating PH features does not explicitly capture the relative location of leukocytes to cancer cells. The relationship between cancer cells and leukocytes is critical as the spatial exclusion of leukocytes from cancer cells provides a mechanism for cancer cells to evade immune surveillance and is observed in many cancer types, including lung adenocarcinoma. To explore the spatial relationship of cancer cells and leukocytes, we computed Dowker PH features between cancer cells and leukocytes (Fig. 1D, Fig. 5A). When computing Dowker PH, the simplicial complex is built on one class of data points, and the construction of the simplicial complex is contingent on the proximity to data points from the second class (see *Methods*). In practice, this means that only cancer cell organisation near leukocytes is captured in the persistence diagram.

**Figure 5.**
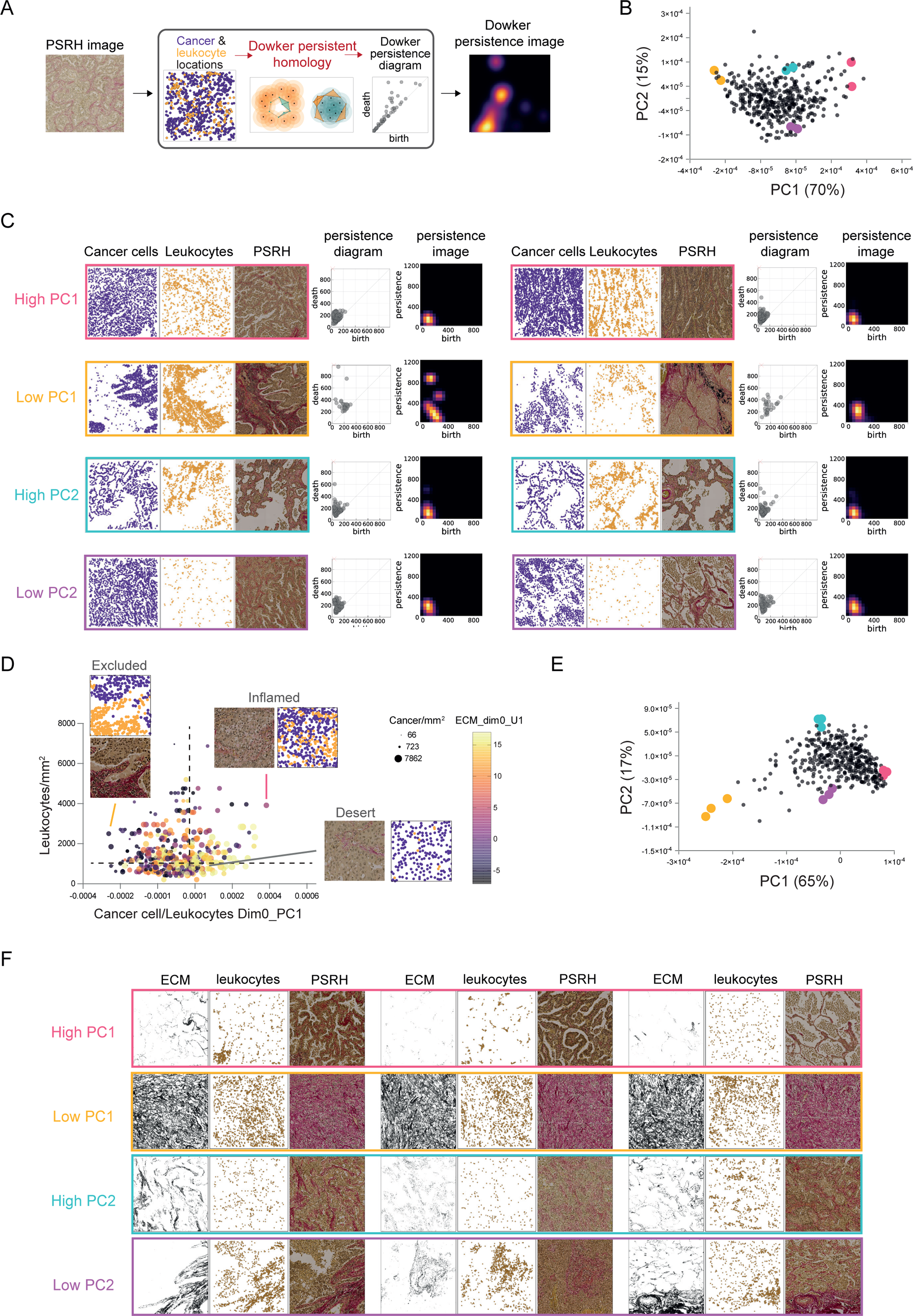
Dowker PH features capture immune exclusion and reveal links to ECM patterns. **A**. Pipeline for computing Dowker PH features. Given an ROI, we computed 0-dimensional Dowker PH features from the locations of cancer cells and leukocytes. The persistence diagram was vectorized into a persistence image of size 20 by 20, which was then flattened into a vector of size 400. **B.** PCA on the 0-dimensional Dowker PH features between cancer cells and leukocytes. **C.** Example ROIs with high and low PC1 (first principal component) and PC2 (second principal component) coordinates. A comparison of example ROIs with high and low PC1 coordinates indicate that PC1 encodes exclusion between cancer cells and leukocytes. In particular, the persistence image for ROIs in with low PC1 coordinates have small values (black) near the origin. A comparison of example ROIs with high and low PC2 coordinates suggest that PC2 can capture colocalizations between cancer cells and leukocytes. **D.** Plot shows PC1 coordinates from panel B (x-axis), leukocyte density (y-axis), UMAP component 1 of 0-dimensional PH features from ECM (colour), and cancer cell density (dot size). Three representative ROIs are shown alongside the position of cancer cells (purple) and leukocytes (yellow). Together, these features enable the distinction of regions that are lacking leukocytes (desert), replete with leukocytes but excluded from cancer cells (excluded), or replete with leukocytes interspersed with cancer cells (inflamed). **E.** PCA on 0-dimensional Dowker PH features on ECM and leukocytes. **F.** Example ROIs with high and low PC1 and PC2 coordinates in panel E suggest that the principal components encode abundance (PC1) and colocalization (PC2) of ECM and leukocytes. Each image is 878µm x 878µm.

We then performed PCA on the 0-dimensional Dowker PH features between cancer cells and leukocytes (Supp. Fig. 7A shows the weighting of different regions of the persistence images for PC1-4 of 0-dimensional Dowker PH features between cancer cells and leukocytes). PCA revealed considerable diversity in the inter-relationship of cancer cell and leukocyte positions, with ROIs with spatially distinct and mutually exclusive localisation of the two cell types reflected in the first principal component (Fig. 5B&C). Low PC1 coordinates reflected clear immune exclusion, while high PC1 coordinates reflected mixing of cancer cells and leukocytes. Intermediate values were associated with immune exclusion at shorter length scales.

We next investigated the relations between 0-dimensional Dowker PH features and other features, such as ECM PH features, cancer density, and leukocyte density. The combination of 0-dimensional Dowker PH features with cancer and leukocyte density enabled the categorisation of ROIs into the three paradigmatic TME classes – inflamed/hot, excluded, and desert. Fig. 5D shows the relationship between PC1 from Fig. 5B (x-axis), which reflects separation of cancer and immune cells over longer length scales), and leukocyte density (y-axis). Furthermore, it additionally reflects cancer cell density (dot size), and UMAP1 from 0-dimensional PH features of ECM in Fig. 2B (dot colour), which reflects ECM abundance. ROIs with low leukocyte densities correspond to ‘immune desert’ regions. ROIs with higher leukocyte numbers can be divided into ‘inflamed/hot’ and ‘excluded’ based on the PC1 coordinate from Fig. 5B. Interestingly, PC1 is associated with distinct patterns of ECM organisation (note how the shading reflecting ECM organisation varies with PC1 values in Fig. 5D), potentially supporting an instructive role of the ECM in the relative spatial positioning of cancer cells and leukocytes. More detailed analysis indicated that spatial segregation of cancer cells and leukocytes was correlated with smaller gaps, altered gap shapes, lower fractal dimension, and altered long range texture features (Supp. Fig. 7C). No clear associations were observed with tumour stage or histological growth pattern (Supp. Fig. 7D).

Dowker PH features can be computed on any pair of the TME constituents. There are thus six distinct Dowker PH features: the 0-dimensional and 1-dimensional Dowker PH features from (cancer, leukocytes), (cancer, ECM), and (ECM, leukocytes). Fig. 5E&F show PCA analysis of the 0-dimensional Dowker PH feature of ECM and leukocytes; interestingly, high PC2 values identified regions with co-localised clusters of leukocytes and ECM. These analyses illustrate the ulitility of Dowker PH for describing the spatial inter-relationships of different TME features. A correlation matrix summarising the relationships between all numerically quantifiable spatial (both PH and Dowker PH) and clinical features is shown in Supp. Fig. 8.

### Linkage of spatial patterns to transcriptomic data

Both PH and Dowker PH features provide powerful descriptors of tissue organisation and reliably identify tumour regions with similar organisation. However, they do not provide insight into the possible mechanisms underpinning the different spatial patterns in lung adenocarcinoma. To obtain these insights we related the PH features and their clusters to other classes of data about the tumours, in particular, to data from transcriptomic analysis. The ROIs from the whole slide images used for the analysis presented thus far lack region-specific transcriptional analysis. Fortunately, the TRACERx lung study includes regional biopsies that have been analysed by RNA sequencing and whole-exome sequencing [30–32] (see *Methods*). Cored tissue samples from multiple tumours and tumour regions were constructed into a tissue microarray (TMA). We stained the TMAs using the same PSRH protocol applied to the diagnostic blocks (see *Overview of pipeline for extracting topological features of lung adenocarcinoma*). 71 PSRH images from 44 tumour regions (22 patients) were taken forward for analysis. We computed the PH features, Dowker PH features, and cell type density metrics from our machine learning method (see *Methods*), and, for the 29 tumour regions with paired RNA sequencing data, linked these imaging features to transcriptional data.

We focussed on determining transcriptional programmes linked to features of ECM architecture. Fig. 6A shows PCA of 1-dimensional PH features on ECM for 62 samples for which ECM PH features were calculated (derived from 38 tumour regions, 20 patients). The points are coloured according to the closest clusters in Fig. 2B. We observed similar transitions among clusters as the transitions in Fig. 2B, with increasing ECM density reflected in PC1 and ECM texture in PC2. We categorised the tumour regions into those with values above or below the median for each PC in turn. We then performed gene set enrichment analysis on the subset of samples for which paired RNA sequencing data were available to interrogate Hallmark pathways [42] enriched in regions characterised as having an above-median versus below-median PC value and vice-versa (see *Methods*). Of note, the dimension corresponding to increasing ECM density (PC1) is associated with metabolic changes, including a downregulation of genes linked to oxidative phosphorylation, fatty acid metabolism, and reactive oxygen species. To expand on our analysis of individual principal components, we also compared gene expression patterns among ROIs assigned to different ECM organisation clusters (41 samples, 25 tumour regions, 17 patients) (see *Methods*). For this transcriptomics analysis, we further grouped clusters together based on the pseudotime analysis in Fig. 2F to ensure that each group being compared had sufficient samples within it (five or more). Interestingly, this identified inflammatory signalling and epithelial to mesenchymal transition as higher in the partial fibrosis end-stage defined by clusters 4 and 5 compared to other clusters, including the fully fibrotic end-stage (Fig. 6B). Thus, PH features enabled transcriptional correlates of spatial ECM patterns to be discovered.

**Figure 6.**
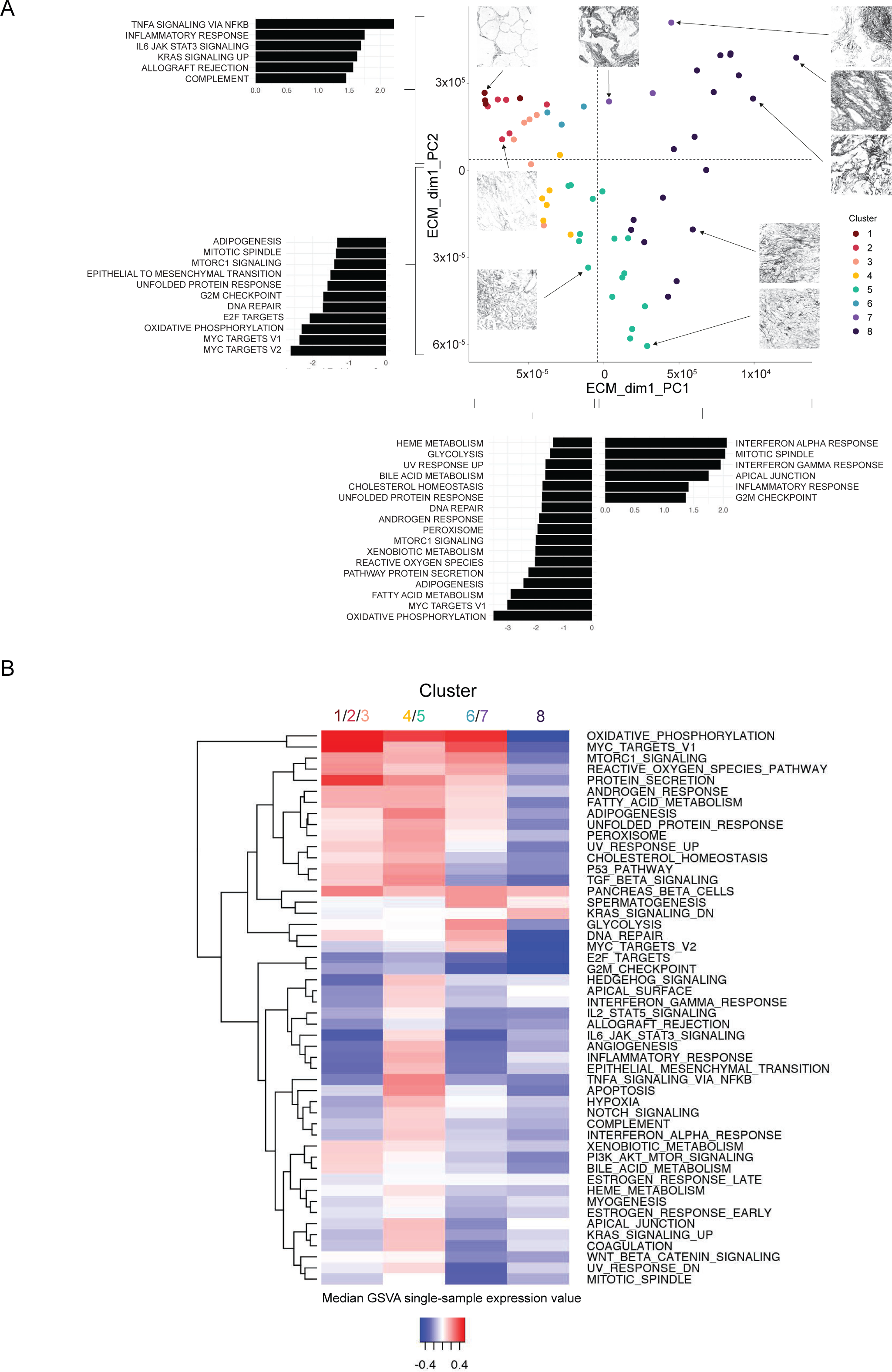
Linking topological features to transcriptional programmes. **A.** PCA of 1-dimensional PH features of ECM of 62 tissue microarray-derived lung adenocarcinoma samples (38 tumour regions, 20 patients). Colours indicate the ECM cluster (as defined in Figure 2) to which each sample belongs. Plot is divided into quadrants based on the median values for PC1 and PC2, and genesets enriched in samples assigned as *High* compared to *Low* or *Low* compared to *High* PC1 or PC2 values are shown alongside the plot accordingly, with the length of the bar reflecting the normalised enrichment score. Numbers of tumour regions contributing to GSEA analysis - PC1: *High* 11 tumour regions, 10 patients; *Low* 11 tumour regions, 7 patients; PC2: *High* 14 tumour regions, 10 patients; *Low* 9 tumour regions 8 patients. **B.** Heatmap shows the differential expression of the indicated Hallmark genesets [42] in ECM clusters 1-3 (reflecting early stages of ECM changes), clusters 4-5 (reflecting one of the end stages), clusters 6-7 (reflecting later intermediate stages), and cluster 8 (reflecting almost fully fibrotic tissue), performed using single-sample gene set variation analysis and ECM clusters as defined in Figure 2. Analysis performed for 41 samples, 25 tumour regions, 17 patients (clusters 1-3: 14 samples, 9 tumour regions, 6 patients; clusters 4-5: 10 samples, 9 tumour regions, 9 patients; clusters 6-7: 5 samples, 5 tumour regions, 5 patients; cluster 8: 12 samples, 9 tumour regions, 8 patients).

### An integrated understanding of spatial changes in the tumour microenvironment

Lastly, we sought to combine information from the PH features and pairwise Dowker PH features to provide an integrated perspective on spatial patterns in lung adenocarcinoma. To this end, we generated a UMAP after concatenating all the PH features and Dowker PH feature vectors (Supp. Fig. 9A). Once again, the analysis confirmed the ability of topological features to group tumour regions with similar spatial patterns of tumour microenvironment organisation, with individual clusters having interpretable features. We next returned to the pseudo-time analysis of 0-dimensional and 1-dimensional PH features of ECM and overlaid other features on the inferred temporal sequence. This revealed that the density, branch points, and fractal dimension of the ECM increased with pseudo-time, while the size of gaps reduced (Supp. Fig. 9B). These changes were more pronounced in the pseudo-time end stage containing ECM clusters 8&9, which reflects almost total tissue fibrosis. Increasing pseudo-time was also accompanied by increasing leukocytic infiltrate and decreasing numbers of cancer cells (Fig. 7A). More notably, the cancer cell – leukocyte Dowker feature (PC1 from 0-dimensional Dowker PH feature between cancer cells and leukocytes) decreased as pseudo-time increased (Fig. 7A). This phenomenon is reflective of spatially exclusive patterns of cancer cells, indicating that immune exclusion increases with time (CC-L Dim0_PC1 in Fig. 7A). Interestingly, immune exclusion was less pronounced in the end-stage reflecting partial fibrosis. Consistent with this, the ECM – leukocyte feature reflecting immune cell localisation away from ECM-rich areas (ECM-L Dim0_PC2in Fig. 7A) was high in the partial fibrosis end-stage, but low in the fully fibrotic end-stage. This indicates that not all features undergo progressive and linear changes over time. Collectively, our analysis supports a model in which cancer initiates in tissue with relatively normal ECM organisation, although some thickening of the ECM is observed in non-cancerous tissue and may precede cancer initiation. There are then progressive changes, with increasing ECM density, fractal dimension, and reducing gap sizes. These can lead to two different end states: one with elevated ECM, but with leukocytes not restricted to ECM areas, and the other reflecting a highly fibrotic state, with few cancer cells and high levels of immune exclusion.

**Figure 7.**
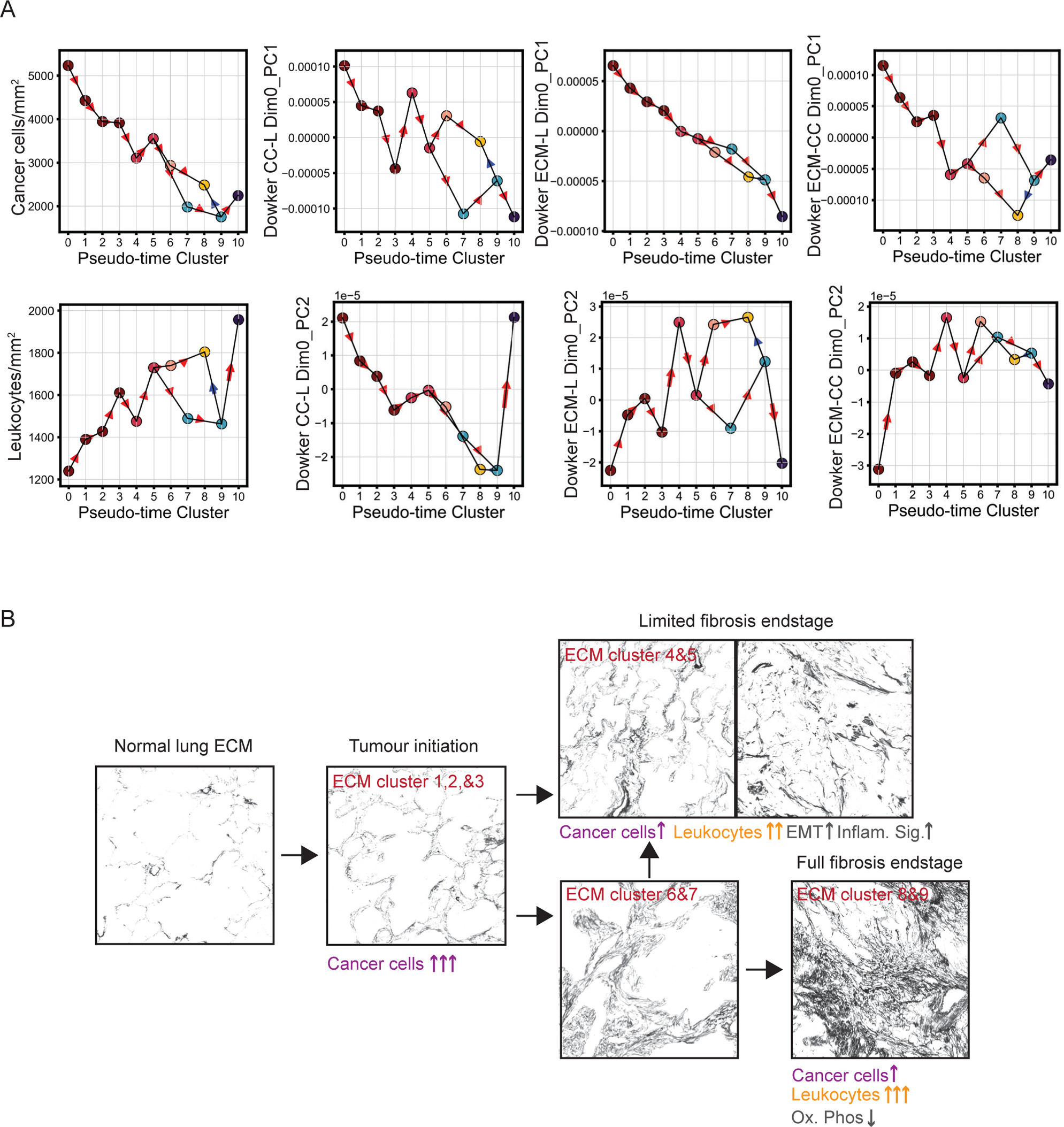
Changes in the TME that accompany progressive ECM remodelling. **A.** Plots show how the indicated features vary as a function of pseudo-time cluster (x-axis). The colour of the dot reflects the predominant ECM cluster (based on Figure 2) at that time point. **B.** Scheme showing information distilled from the preceding figures. Initial ECM organisation is similar to that found in normal lung tissue. The trajectories and changes associated with the two different end stages are shown. Grayscale images show collagen staining extracted from PSRH images. Each image is 878µm x 878µm.

## Discussion

In this study, we quantify spatial patterns within lung adenocarcinomas so that similar tumour microenvironments can be identified and grouped together, and so that spatial patterns can be related to other features of the disease. Specifically, we focus on analysing the ECM and its relationship to cellular distributions. We employ methods from topological data analysis to describe spatial patterns because they are adept at capturing density and clustering information, which is highly relevant for cells, and quantifying loops and gaps, which is pertinent to ECM organisation. Moreover, by employing Dowker persistent homology, we develop a framework for quantitatively describing the inter-relationship of different TME components across different length scales.

Numerous groups have developed features for the analysis of ECM fibres [43–45] (Fu, Jenkins *et al.*, in preparation). These include quantitative measurements of ECM density, fibre numbers, fibre branching, fibre alignment, fractal feature, texture features, and gaps. These features have provided useful insights into how ECM structure changes as disease progresses, and they have been linked with patient outcomes [10, 46]. However, these methods are not well suited to incorporating information about cellular position, although this can be done via post-hoc spatial statistics. Other studies have sought to relate cell position to tumour stroma boundaries, usually via simple distance metrics [47]. These are valuable, but by reducing to a one-dimensional metric, information about the broader spatial organisation of the cells and tumour stroma boundary is lost. Tumour topological graphs are better able to describe longer range organisation of cells as they build local graphs of cell positions and can determine the frequency with which two cells have a third cell type between them [13]. However, this type of analysis has not been applied to ECM, partly because ECM is a continuous material that is not readily discretised in the way that cells are. By treating both cell positions and ECM as point clouds and with attention paid to sampling – specifically greater sampling of the ECM to reflect its more continuous nature – we use topological methods to analyse the relative position of different cell types and the ECM in cancer tissue. Excitingly, Dowker PH captures the spatial relationship between two cells types, or between one cell type and the ECM. This is of utility for the analysis of tumour-immune surveillance, which involves leukocytes engaging with cancer cells. High levels of leukocyte infiltration, particularly of CD8 T-cells, are a favourable prognostic indication in many cancer types [48–53]. In contrast, either an absence of leukocytes or their location in compartments at distance from cancer cells is unfavourable. We show that Dowker PH can quantitatively capture these classes of tissue organisation. In the future, it will be interesting to use further variations of TDA to quantify the spatial organisation of tumours. Analysis of the vectorised persistence images can be tailored to probe spatial patterns over length scales of particular interest. Multiparameter persistence examines the spatial features as multiple parameters vary, and has been deployed to relate hypoxic regions to the distribution of immune cells [38]. Systems consisting of more than two features of interest can be interrogated using the recently-developed tools of multispecies Witness filtration [28] and Chromatic Alpha complex [54].

Although changes in ECM architecture in cancer have been described by many researchers [2–9], it is less clear in what order these changes might occur. This is partly due to the impracticality of taking repeated biopsies from patients. By sampling a large number of regions spanning diverse ECM patterns and utilizing topological methods to capture patterns across different length scales, we are able to derive a rich dataset of metrics for ECM architecture. The high dimensional nature of the data is suitable for methods that infer likely routes of transition between different ECM states. By applying such methods to our data, we derive likely trajectories of transitions in ECM organisation. This begins with normal lung organisation and is associated with increasing ECM deposition. Interestingly, we observe some similar changes in adjacent non-cancerous lung tissue which indicates that some modest changes in ECM organisation may precede tumour initiation. Moreover, our analysis of neighbouring ROIs suggest that ECM changes propagate laterally in tissue. Our analysis also indicates two different end states; one reflects extensive fibrosis while the other retains ECM gaps. The multi-faceted nature of our analysis enables us to track other changes that accompany altered ECM architecture. ECM patterns were not simply reflective of either tumour stage or histological sub-type. This likely indicates that structural changes in the stroma are not tightly coupled to cancer cell growth. Transcriptional analysis suggests a down regulation of oxidative metabolism and an increase in some inflammatory signalling. Consistent with this, leukocyte numbers increase along the ECM pseudo-time trajectory; however, this is also accompanied by a decrease in PC1 of Dowker PH features between cancer cells and leukocytes, which reflects an increase in immune exclusion. This argues that the progressive changes in ECM that occur in tumours protect the cancer cells from immune surveillance.

The focus of this study was to understand how different ECM patterns relate to each other and to the distribution of different cell types. The limited cohort size (44 patients) precluded analysis of association between topological features and patient outcomes. The future application of TDA to large patient cohorts with clinical follow up will enable spatial patterns associated with disease progression or recurrence to be identified. Given the concept of immune exclusion preventing immunotherapy efficacy [55], it will be interesting to probe how the relationship between distinct immune cell types and the ECM impacts on the response to immunotherapies. As we demonstrate, this could be measured using Dowker PH. Additionally, the relationship between CD8 T-cells and immune-suppressive myeloid cells could be queried by the same methodology.

To conclude, we demonstrate that TDA, namely PH and Dowker PH, is a powerful tool for describing and comparing spatial patterns. Moreover, the resulting high dimensional descriptions of spatial pattern can be interrogated using the same methods as other high dimensional data, including UMAP, PCA, and pseudo-time analysis. Using the latter method, we show how the ECM architecture changes progressively over time. It is likely that ECM patterns are particularly suited to this analysis as they change over longer timescales than cell positions, especially leukocytes that can move at speeds of several microns per minute [56]. By cross-referencing the progression of ECM patterns with other data for the same regions of tissue, we are able to construct a holistic view of how the tumour microenvironment evolves over time in lung adenocarcinoma. We believe that our approach has the power to transform our understanding of how spatial patterns develop over time in pathological conditions.

## Methods

### Data analysed in this study

The data from this study are those for a subset of the first 421 patients prospectively recruited to and with tumour profiling undertaken for the TRACERx study (TRAcking non-small cell lung Cancer (NSCLC) Evolution through therapy (Rx); https://clinicaltrials.gov/ct2/show/NCT01888601, approved by an independent research ethics committee, 13/LO/1546). Informed consent for entry into the TRACERx study was mandatory and obtained from every patient. Methods for data obtention, including the tissue sampling approach during surgery, have been previously described in [30, 31].

#### Tumour diagnostic blocks (whole slide and ROI analyses)

44 whole slides (typically 15mm x 30mm) were obtained from lung adenocarcinoma (LUAD) tumour diagnostic blocks from surgically resected tissue. Details of central histopathological review undertaken on LUAD diagnostic slides to confirm tumour subtype and assign growth pattern information are provided in [35]. Four hundred 878μm x 878μm regions of interest (ROIs) were selected from these whole slide images based on diversity of ECM and cellular organisation. Tumour-level clinical and mutations data used in this manuscript were derived as described in [31]. Tumour mutational burden (mutations/megabase) in Supp. Figs. 4 and 8 was calculated using a harmonized approach [57] using the TRACERx mutation WES calls from [31].

#### Tumour region-level samples (**Figure 6**)

Snap frozen multi-region tumour samples within a surgical resection specimen were processed to formalin fixed paraffin embedded (FFPE) blocks after first taking sufficient material for DNA and RNA sequencing. A single representative core was taken from each regional FFPE block (1.5mm diameter) and cores arranged into tissue microarrays. 71 lung adenocarcinoma cores (71 samples, 44 tumour regions, 22 patients) were taken forward for TDA analysis. Region-level RNA data were available for 29 of these tumour regions from 18 patients and were processed and gene expression values derived as described in [32].

### PSRH staining

Samples were stained using a combination of Weigert’s iron haematoxylin solution (Sigma HT1079) and a Picrosirius Red Kit (Abcam ab150681). Briefly, slides were deparaffinised and hydrated, before incubation in Weigert’s Haematoxylin solution for 10 minutes and then in Picrosirius red solution for 60 minutes, rinsing twice in acetic acid, then alcohol dehydration, and finally mounting. Slides were then scanned at 20× using Zeiss Axio Scan Z1.

### Identification of cell types in sections

#### 1. Dataset preparation

The dataset for training and validating the cell identification model was collected from 10 picrosirius-red-stained whole slide images (WSI) of lung adenocarcinoma. We denoted regions of interest on the WSI and dot-annotated every single cell identifiable within the regions. Cells were assigned one of five types based on their morphology; the five cell types correspond to cancer cells, normal epithelial cells, leukocytes, fibroblasts, and blood cells. We also labelled necrosis, black pigments, and other cell types without determinable identities. The 10 slides were randomly split into training and testing datasets with an 8:2 ratio. For the 8 slides of the training dataset, regions containing annotations were scaled to a resolution of 0.44 μm/pixel and were then cropped into 224×224 patches with a stride of 120 pixels. Patches were carefully inspected to ensure every cell and tissue element identifiable on the patch were labelled, which gave rise to 12240 annotations on 286 patches in total. For the testing dataset, we collected 1213 annotations from 2 WSIs. A breakdown of annotations for each category in the training and testing datasets is shown below.

| Cell types | Training | Testing |
| --- | --- | --- |
| Cancer/Epithelial | 5283 | 567 |
| Leukocytes | 5039 | 198 |
| Necrosis | 390 | 31 |
| Fibroblast | 792 | 139 |
| Pollution | 346 | 72 |
| Red blood cells | 246 | 133 |
| Others | 144 | 73 |
| **Total** | **12240** | **1213** |

#### 2. Model training and testing

We used a double-branch model S-DRDIN [58] to predict element locations and classes simultaneously. The model has been shown to outcompete single-task models by benefiting from the integration of cell detection and classification features [58]. To train the model, we converted dot annotations on 224×224 patches into classification masks of nine channels, with each channel representing locations of one category, and the last channel being a background mask representing locations without any element present. We prepared another detection mask per patch denoting all elements identified on the patch without categorical information. Data augmentation of horizontal and vertical flipping was performed. The model was trained with both detection and classification masks as inputs. We trained the model for 100 epochs with a batch size of 4. Weights were initialized using uniform glorot [59] and optimized using Adam [60] with a learning rate of 0.001.

The trained model was applied to regions of interest to generate a probability map of cell locations. The map was binarised by a threshold of 0.05, followed by the identification of connected components. Components smaller than 60 pixels were discarded to reduce false positive detection. The predicted locations of elements were determined by the local maxima within a sliding window of size 15×15 pixels. We then averaged the predicted values of each category in a 49×49 pixel square centred at the detected element. An element was assigned the category with the highest average probability. For elements predicted as others, the second most probable category was assigned. Cells predicted as cancer cells or normal epithelial cells were merged into one single epithelial group.

The derived cell predictions were tested against manual annotations in the testing dataset. A predicted cell was regarded as a true positive if it was located within 15 pixels of a manually annotated cell of the same category. In general, S-DRDIN achieved an accuracy of 0.864 and a weighted F1 score of 0.860 for six categories, with the F1 score being 0.935, 0.623 and 0.747 for epithelial cells, fibroblasts, and leukocytes respectively – shown below.

| Cell types | Precision | Recall | F1 score |
| --- | --- | --- | --- |
| Cancer/Epithelial | 0.903 | 0.969 | 0.935 |
| Fibroblast | 0.618 | 0.627 | 0.623 |
| Leukocytes | 0.837 | 0.674 | 0.747 |
| Necrosis | 0.538 | 0.467 | 0.500 |
| Red blood cells | 0.917 | 0.901 | 0.909 |
| Pollution | 0.937 | 0.983 | 0.959 |

### Topological data analysis (TDA)

TDA uses tools from topology to study the shape and structure of data.

#### Simplicial complexes and homology

A simplicial complex is a generalization of a graph that consists of vertices, edges, triangles, and their higher-dimensional analogues. Given point cloud data, one popular way of constructing a simplicial complex is to use the distances among the data points as follows. Fix some distance parameter. Take the data as the vertex set. For each pair of points whose distance is at most *ε*, create an edge between 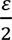 the corresponding vertices. Equivalently, one can draw balls of radius around each point and create an edge between vertices when the balls intersect. For each tuple of points, if all the pairwise distances among the three points are at most *ε*, then create a triangle among the three corresponding vertices. If (*n* + 1) points have pairwise distances at most *ε*, then we add a higher-dimensional version of a triangle, called an *n*-simplex (Fig. 1B). Let *X* denote the resulting simplicial complex.

Properties of *X*, such as connected components, loops, and voids, are computed via homology. The 0-dimensional homology of *X*, denoted *H*_0_(*X*), is a vector space whose dimension equals the number of connected components in *X*. The 1-dimensional homology, denoted *H*_1_(*X*), is a vector space whose dimension equals the number of loops in *X*. The *n*-dimensional homology, denoted *H*_*n*_(*X*), is the vector space whose dimension equals the number of -dimensional features in *X*.

#### Persistent homology

Given a point cloud, we first create a sequence of simplicial complexes

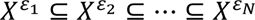

where *X*^*εi*^ is the simplicial complex with distance parameter *ε*_*i*_ as constructed above. Persistent homology summarizes the birth and death of topological features (*H*_0_ and *H*_1_) across the sequence. To study the birth and death of *n*-dimensional topological features, we examine the changes in the *n*-dimensional homology *H*_*n*_ across the sequence. A feature that is born at parameter *b* and dies at parameter *d* is represented by a point with coordinates (*b*, *d*). The collection of such points is referred to as the *n*-dimensional persistence diagram. The evolution of clusters is summarized in a 0-dimensional persistence diagram. A point (*b*, *d*) indicates that a cluster is born at parameter and dies at parameter by merging with another cluster. The evolution of loops is summarized in a 1-dimensional persistence diagram. A point (*b*, *d*) indicates that a loop forms at parameter *b* and that this loop is filled-in by a collection of triangles at parameter *d* (Fig. 1B).

The points on a persistence diagram with a large death parameter and a small birth parameter, which occupy the region that is far from the diagonal line, are usually considered significant features while points that are closer to the diagonal are considered noise.

#### Dowker persistent homology

Dowker persistent homology [24, 25] allows the study of structural relations between pairs of point clouds. Let *P* and *Q* be two point clouds. Given a distance parameter *ε*, we first construct a Dowker complex 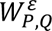 that has *P* as the potential vertex set. We add an edge (a 1-simplex) between points *p*, *p*_1_ ∈ *P* if there exists a point *q* ∈ *Q* whose distances to *p*_0_ and *p*_1_ are both at most *ε*. We add a triangle (a 2-simplex) among three points *p*_0_, *p*_1_, *p*_2_ ∈ *P* if there exists a point *q* ∈ *Q* whose distances to *p*_0_, *p*_1_, *p*_2_ are at most *ε*. Similarly, we add an *n*-simplex [*p*_0_, …, *p*_*n*_] if there exists a point *q* ∈ *Q* whose distances to *p*_0_, …, *p*_*n*_ are at most *ε*. We then create a nested sequence of Dowker complexes for varying distance parameter

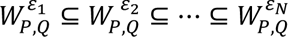

We again study the evolution of *n*-dimensional topological features by examining the change in homology *H*_*n*_ across the sequence. The birth and death of -dimensional features are summarized in a *n*-dimensional persistence diagram. To distinguish from the standard persistence diagram, we refer to the result as *n*-dimensional Dowker persistence diagram. By Dowker’s Theorem [24, 25] the Dowker persistence diagram that results from the above sequence (with *P* as the vertex set) is identical to the Dowker persistence diagram that results from the following sequence that uses *Q* as the vertex set:

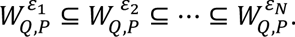

#### Vectorizations of topological descriptors

We vectorize (Dowker) persistence diagrams using persistence images [61] Given a (Dowker) persistence diagram *PD*, consider *PD* as a collection of birth-death coordinates. We first transform *PD* by mapping each point (birth, death) to (birth, death − birth). Let *T(PD)* denote the transformed persistence diagram. We then create a surface by taking a weighted sum of Gaussian distributions centered at each of *T(PD)*. The Gaussians are weighed proportional to death − birth of the center. The resulting surface is discretized into an array. The 0-dimensional persistence diagram is discretized to a vector of length 20, and the 1-dimensional persistence diagram is discretized to an array of size 20 by 20, which we refer to as the persistence image (PI). The 1-dimensional PI is flattened into a vector of size 400 prior to analysis. See Supp. Fig. 1D for a visualization of the vectorization process.

### Analysis of tissue microarray data and linkage to transcriptional profiles

71 TMA-embedded LUAD tumour sections were stained with PSRH. Each section represented tissue from a single tumour region acquired by TRACERx multi-region sampling, as described in [31]. 44 distinct tumour regions from 22 patients were represented in the 71 sections (>1 tumour region n=13 patients, 1 tumour region n=9 patients). TDA features were paired with RNAseq data generated from paired region-level samples [32] to analyse the relationship between TDA and transcriptomic features. 27 tumour regions were represented twice in the TMAs, and statistical analyses were adapted to adjust for these duplicates as described below.

TDA features were extracted for ECM for 62 of the 71 samples using the pipeline already described. The first two principal components were extracted from PCA of the dimension 1 PH features of ECM data (PC1, PC2) and samples assigned as either *High* or *Low* for each principal component in turn based on splitting each principal component by its respective median. Any tumour region represented by two tissue samples for which one tissue sample had a *High* coordinate value and the other *Low* was excluded from analysis to prevent identical transcriptomic data contributing to both the High and Low category for a single principal component.

Data remaining after these inclusions/exclusions comprised: PC1: *High* 11 tumour regions, 10 patients; *Low* 11 tumour regions, 7 patients; PC2: *High* 14 tumour regions, 10 patients; *Low* 9 tumour regions 8 patients. Differential Gene Expression Analysis (DGEA) was run on the remaining data between *High* and *Low* categories in the following way, as detailed in [32]. First, trimmed mean of M-values normalization from the edgeR (v.3.26.5)[62] R package was performed on RNA-Seq by Expectation-Maximization (RSEM) raw counts. Genes with expression <30 counts/million in <70% of the smallest group size were removed using the function filterByExpr() with min.count set to 30. Expression differences were performed at the region level through the limma-voom analytical pipeline, taking tumour as a blocking factor, by performing within-tumour expression correlations and including them within the voom model estimate using the duplicateCorrelation() function. Following DGEA, gene-set enrichment analysis (GSEA) was run using the R package fgsea (v.1.10.1)[63] using the t-statistic generated by limma as input and with default parameters to identify individual MSigDB Hallmark Gene Sets (v2023.1) (https://www.gsea-msigdb.org/gsea/msigdb/human/collections.jsp#H) [42] which were significantly up- or down-regulated between *High* and *Low* levels of each coordinate value (p_adj_<0.05).

For the comparison of Hallmark Gene Set expression by ECM cluster, gene set variation analysis was performed on RSEM-TPM values for the 29 tumour regions for which RNA sequencing data was available, with RSEM-TPM values calculated as described in [32]. Specifically gene set variation analysis was implemented using the R library GSVA (v1.44.5) [64], invoking the gsva function (method=“gsva”,min.sz=5, max.sz=500, kcdf=“Poisson”, mx.diff=TRUE, parallel.sz=1) to derive a single-sample-level geneset enrichment score for each Hallmark Gene Set. The gplots::heatmap.2 function in R (v3.1.3) was then used to plot these enrichment scores for the 41 samples with paired RNA sequencing and ECM cluster information (41 samples, 25 tumour regions, 17 patients).

### Pseudo-time Analysis

For the pseudo-time analysis we used the Python library pyVIA, developed in [65]. The 420 by 1 concatenated ECM PH arrays for each of the sample ROIs was passed directly to pyVIA with true_label being given by our cluster ids for ECM PH in Figure 2B and with the root selected to be Cluster 1. Various choices of pyVIA parameters were explored, with default parameters providing the most robust results. We set random_seed=4 for reproducibility.

### Cluster Neighbourhood Adjacency Analysis

The ROIs from WSI were assigned a closest cluster. For each cluster ID, we identified the ROIs with the given cluster ID and recorded the cluster IDs of their neighbouring ROIs using 4-connectivity. We then calculated the frequency that each pairwise set of clusters neighbour each other. To assess whether these frequencies of cluster neighbours occurred more or less often than chance, we randomly shuffled the cluster IDs over all ROIs while conserving the total number of each cluster ID and computed pairwise neighbouring cluster frequencies. We repeated the random shuffle 1999 times to give a total sample of 2000 including the observed data. For each pairwise cluster neighbour frequency we ranked the data and generated a two-tailed p-value based on where in the ranked list our observed data fell. We also recorded the tail of this significance i.e. whether the observed frequency is above or below expectation from random chance to provide information on whether the clusters are attractive or repulsive. For significance over all WSI, we calculated the median p-value and sign (attractive or repulsive) for each pairwise interaction of cluster neighbours. This approach controls for clusters occupying differing numbers of grid-points.

### Implementation

All code and analysis are available at https://github.com/irishryoon/lung_cancer_TDA. Persistent homology was computed using the Julia packages Eirene [66] and Ripser [67] Dowker persistent homology was computed in Julia using https://github.com/irishryoon/Dowker_persistence. The persistence diagrams were vectorized using https://github.com/mtsch/PersistenceDiagrams.jl.

---

**Supplementary Figure 1 – linked to Figure 1.**
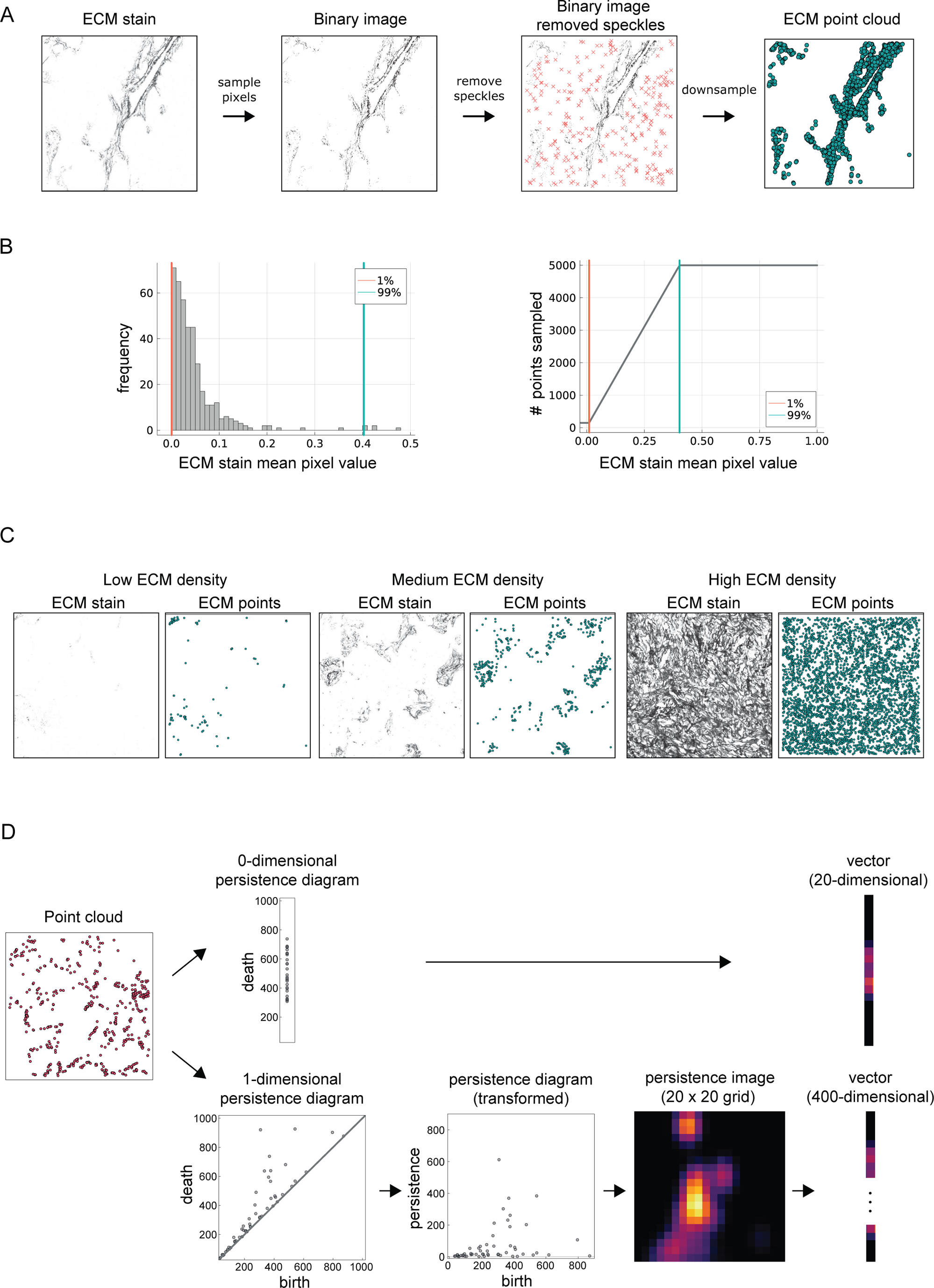
ECM point sampling and vectorisation of a persistence diagram. **A.** An illustration of the ECM point sampling process. Given an ECM stain, we first invert the pixel values and scale them so that the pixel values live in the range [0, 1]. A pixel value of 1 corresponds to black and a pixel value of 0 corresponds to white. For each pixel location, if *p* is the pixel value, we sample a point according to the Bernoulli distribution with a success probability of *p*^2^. This results in a binary image in which the black pixels indicate the points sampled. We then remove points of low density (“speckles”) as follows. For each point *x*, we consider a square region of size 50 pixels by 50 pixels centred at *x*. If such neighbourhood contains less than 5 points, then we remove *x*. Such speckle points are indicated by red crosses. This process removes speckles of high intensity^2^. Lastly, we downsample the remaining points to a target number of points. The target number of points in an ROI is determined according to the mean ECM pixel value as shown in panel B. **B.** Given an ECM stain, the number of ECM points is determined by the stain density. (Left) A histogram of the mean ECM pixel values for the 400 ROIs studied. Most ECM stains have mean pixel values close to zero. The orange and teal vertical lines indicate the bottom and top 1% values. (Right) A function whose input is the mean ECM pixel value of an image and whose output is the number of ECM points to sample. For ECM stains whose mean pixel value is at bottom 1%, we sampled 100 points. For ECM stains whose mean pixel value is within the top 1%, we sampled 5,000 points. For all other ECM stains, the number of points is determined via linear interpolation. **C.** Example ECM stains and sampled ECM points with varying levels of ECM density. **D.** Vectorisation of a persistence diagram. Given a 0-dimensional persistence diagram, we create a weighted sum of Gaussians centered at each point of the persistence diagram and discretize into a vector of length 20. Given a 1-dimensional persistence diagram, denoted *pd*, we first transform *pd* by mapping each point (birth, death) in *pd* to (birth, death - birth). Let *T*(*pd*) denote the transformed diagram. We then create a surface by taking a weighted sum of Gaussians centered at each point of *T*(*pd*). The surface is then discretized into an array of size 20 by 20, which we refer to as a persistence image. Lastly, we flatten the persistence image to obtain a vector of length 400.

**Supplementary Figure 2 – linked to Figure 1.**
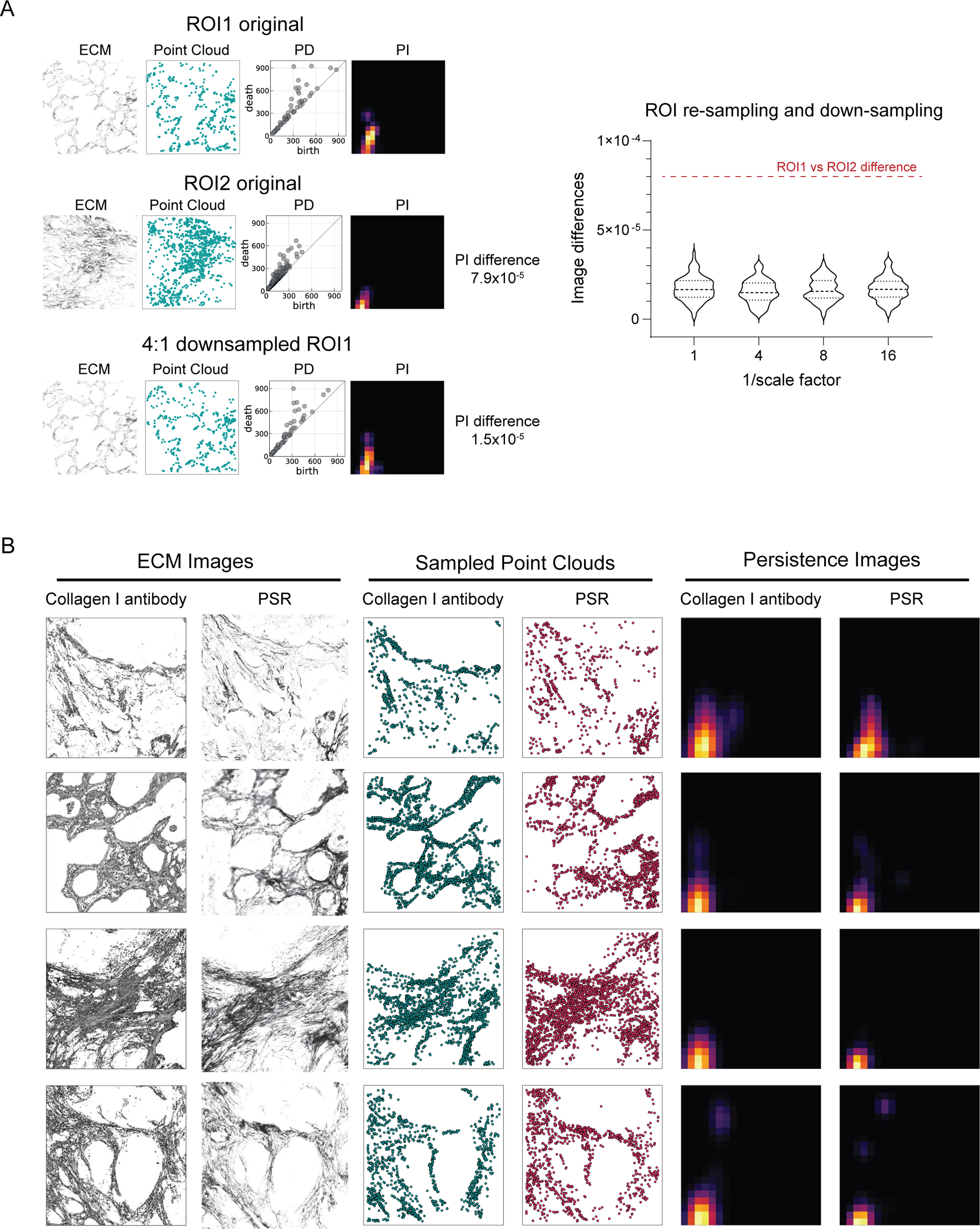
Robustness of topological features to image resolution and acquisition method. **A.** Two ROIs (labelled 1 & 2), their ECM stains, ECM points, 1-dimensional persistence diagrams (PD) and persistence images (PI) are shown on the left. In addition, ROI1 is shown down-sampled 4:1 along with the corresponding point cloud, PD, and PI. For ROI1 and ROI2, the difference in topological features (as measured by the Euclidean distance between the persistence images) is 7.9 x 10^−5^. The right-hand plot illustrates the robustness of topological features. For 100 randomly sampled ROIs (out of 400), we decreased their resolution by factors of 1, 4, 8, and 16. At scale factor 1, we are simply analysing the difference in persistence images that arise from resampling ECM point cloud from the same image. We computed the difference in persistence images and report the box plot of the difference values for each scale factor. Note that the differences consistently stay under 7.9 x 10^−5^, which is the difference in persistence images for ROI1 and ROI2 in panel A. **B.** We show the similarity of topological features (persistence images) obtained from the collagen I stain and the colour deconvolution of PSRH image for 4 exemplar ROIs (one ROI per row). Columns from left to right show the collagen I stain, the colour deconvolution of the PSRH image, the collagen I image-sampled point cloud, the deconvolved PSRH-sampled point cloud, the collagen I-derived persistence image, and the PSRH-derived persistence image.

**Supplementary Figure 3 – linked to Figure 2.**
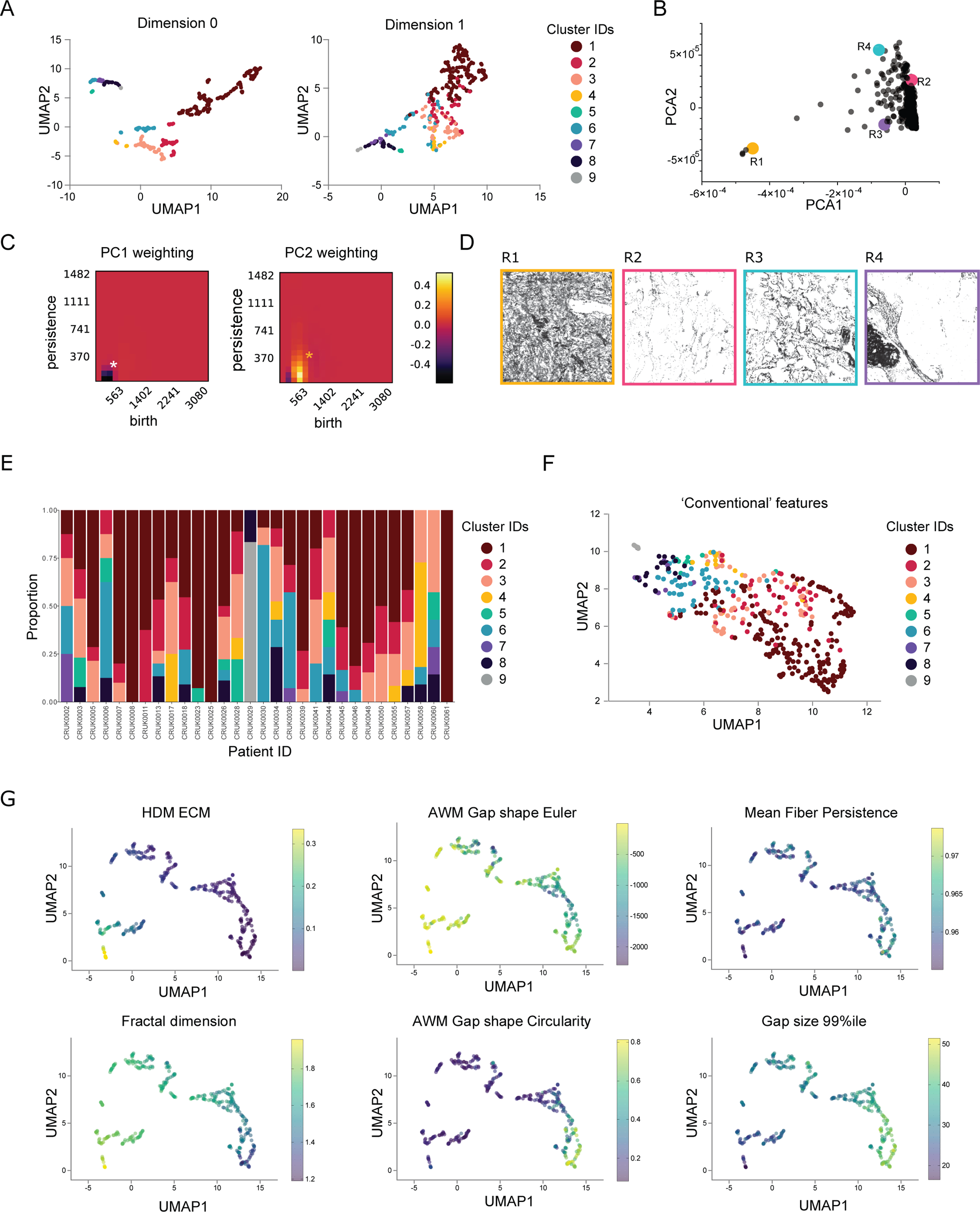
Topological features reveal patterns of ECM architecture. **A.** Applying UMAP on the 0-dimensional PH features from ECM (left) and 1-dimensional PH features from ECM (right) with cluster identities from Fig. 2B overlaid reveals that 0-dimensional PH features effectively separates clusters 1 – 4 and 1-dimensional PH features separates clusters 5 – 8. **B.** PCA on 1-dimensional PH feature vectors of ECM. **C.** A visualization of the first two eigenvectors of the PCA allows interpretation of the principal components in terms of the persistence images. The dark regions indicate a lack of points in the persistence diagram in comparison to the mean, and the bright regions indicate an abundance of points in the persistence diagram in comparison to the mean. **D.** Example ROIs with high and low PC1 and PC2 coordinates. A comparison of R1 and R2 indicates that the PC1 encodes ECM density. A comparison of R3 and R4 indicates that PC2 encodes the presence of high-persistence loops with birth parameter of approximately 350. **E.** Chart shows intra-tumour heterogeneity of patterns of ECM organisation (as defined by clusters in Fig. 2B). Each column represents an individual tumour. **F.** Applying UMAP to features identified using existing spatial tools, including CT-FIRE, gap analysis, and texture features, with colour coding to reflect clusters identified in Fig. 2B. **G.** Features quantified using non-topological tools are represented by varying dot colours on UMAP of ECM PH features in Fig. 2B. Lighter yellow colours indicate higher values. Panels show high density ECM (HDM), average weighted mean (AWM) of gap shape Euler analysis, mean fibre persistence, fractal dimension, AWM of gap shape circularity, and 99^th^ percentile of gap size.

**Supplementary Figure 4 – linked to Figure 2.**
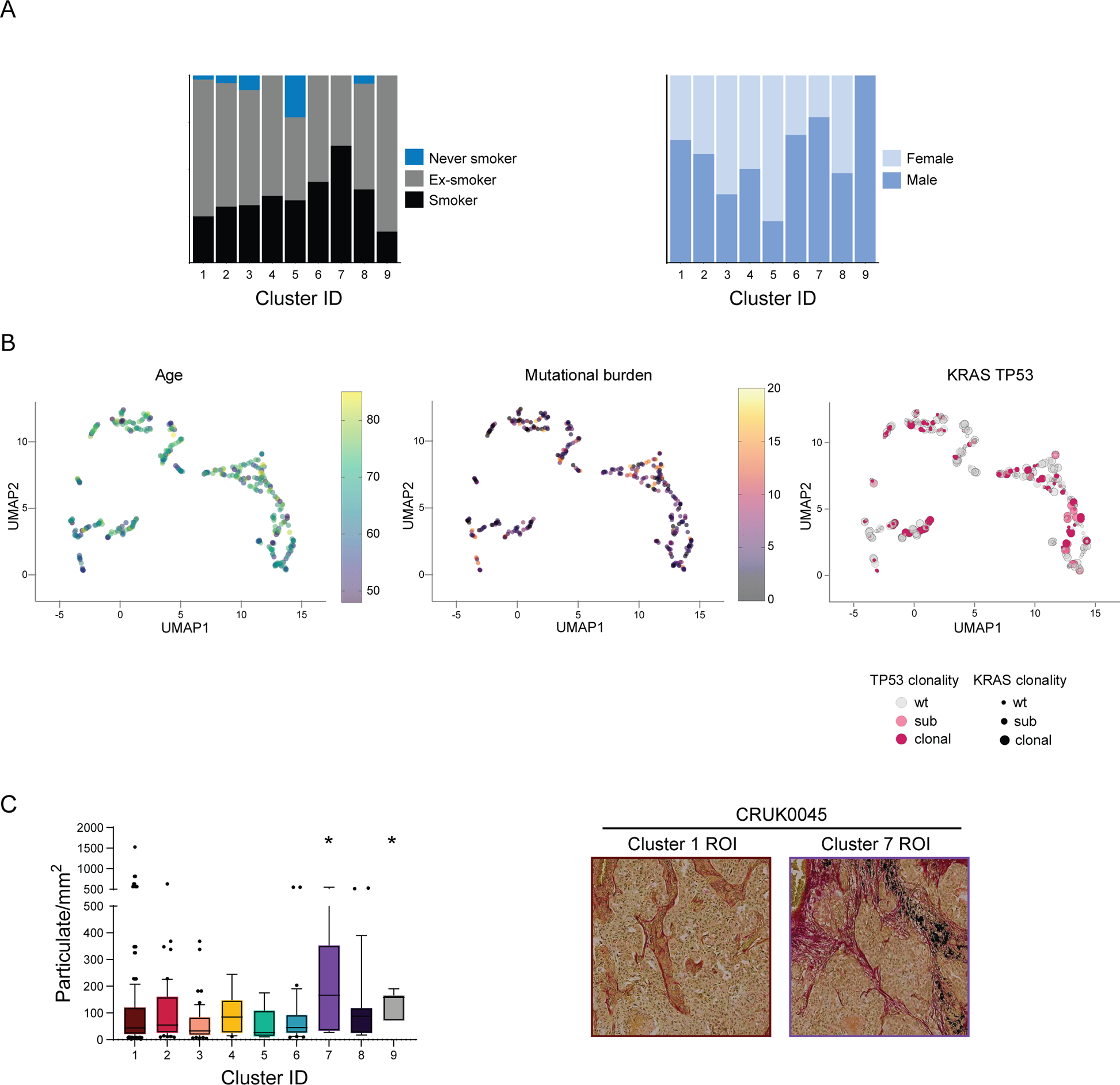
Linkage of ECM PH features to additional clinical parameters. **A.** Charts show a lack of relationship between ECM organisation and (left) smoking status and (right) gender. **B.** UMAP of ECM PH feature is overlaid with (left) age, (middle) tumour mutational burden calculated in mutations/megabase, and (right) KRAS TP53 tumour-level mutation status. **C.** (Left) Plot shows the density of particulate matter, predominantly 2.5 - 10µm in size, in ROI assigned to different ECM clusters. * indicates p<0.05. Clusters 7 and 9 had elevated levels of particulate matter. (Right) Representative PSRH images with high and low particulate levels. Each image is 878 x 878µm.

**Supplementary Figure 5 – linked to Figure 3.**
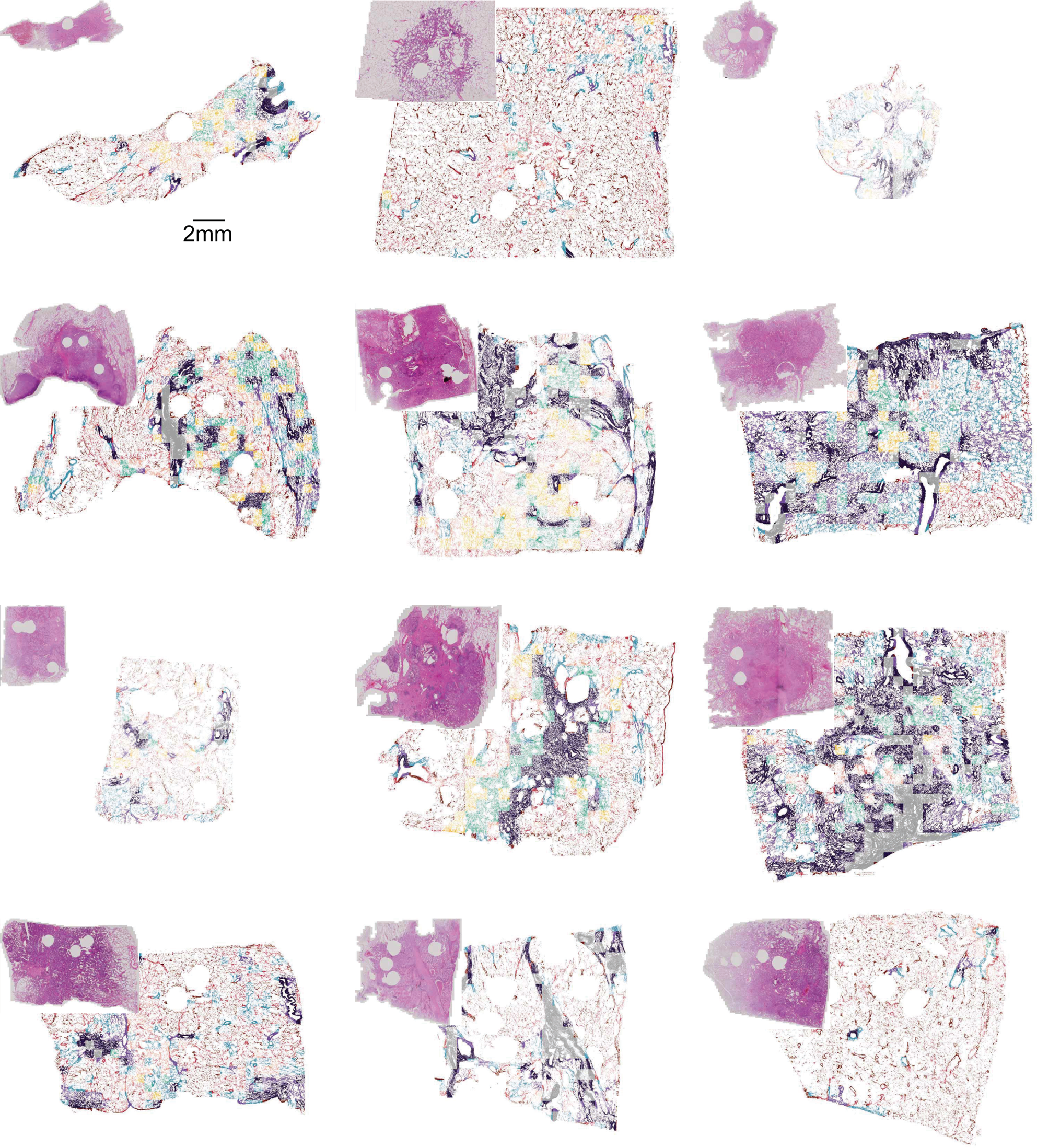
Whole-slide analysis of ECM patterns. **A**. Visualization of a whole-slide tumour with smaller regions coloured by the closest cluster. The whole-slide ECM stain was split into patches of sizes 878μm x 878μm. For each patch, we computed the distance between its topological feature vector and the average feature vector of each cluster (as designated in Fig. 2B). The cluster with the shortest distance was assigned to the patch. The colours indicate the cluster closest to the patch. Inset images show H&E images of a serial section. Scale bar is 2mm.

**Supplementary Figure 6 – linked to Figure 4.**
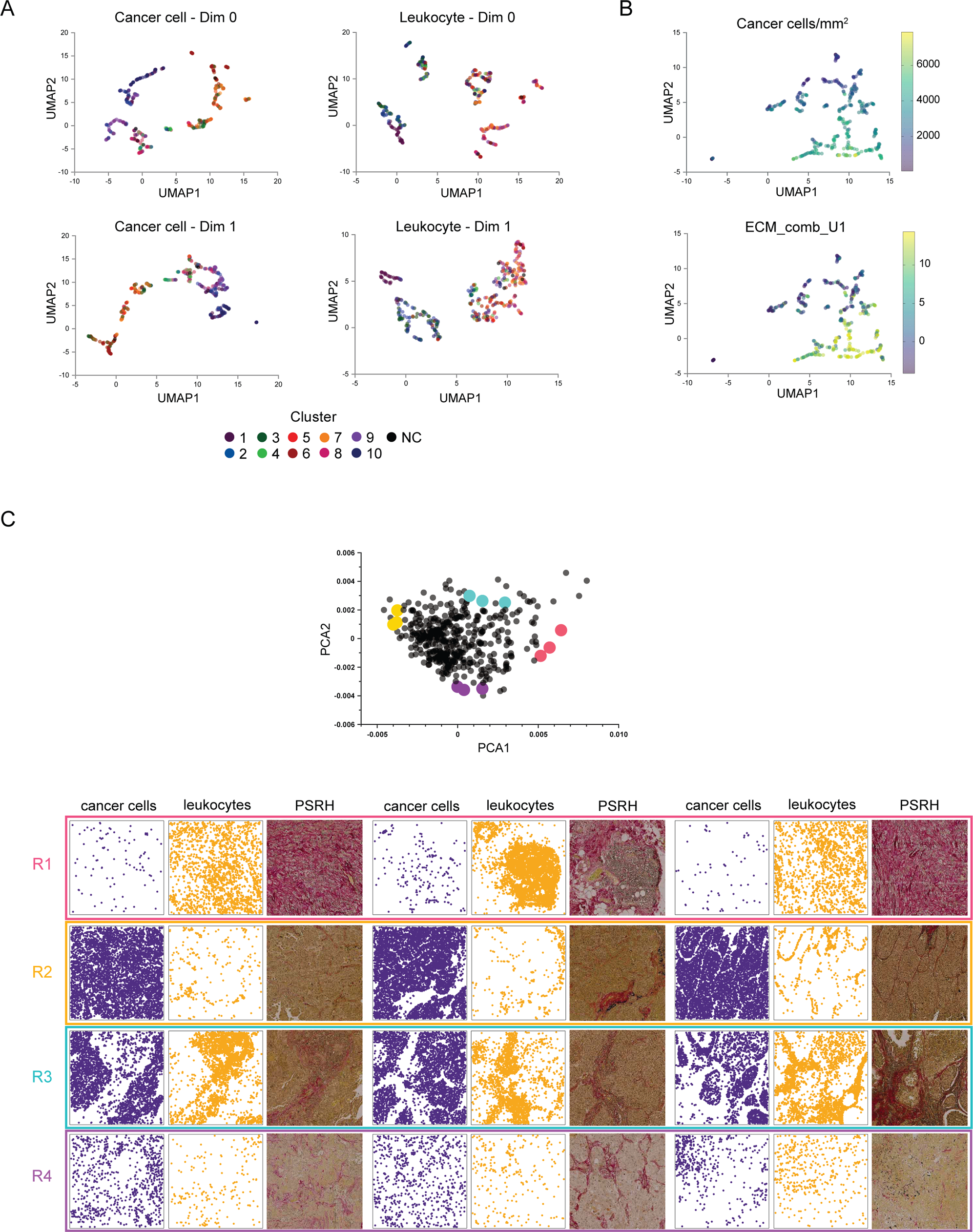
**A**. UMAPs on 0-dimensional and 1-dimensional PH features on cancer cells and leukocytes. Dot colour indicates the cluster assigned to the ROI in the analysis of the combined PH feature in Fig. 4A. NC indicates outlier points not assigned to a cluster. **B**. UMAP on combined PH feature in Fig. 4A coloured according to cancer cell density and UMAP1 coordinate of 1-dimensional PH feature of ECM. The clusters identified by the combined PH feature are correlated with cancer cell density and ECM organisation. **C.** PCA on 0-dimensional and 1-dimensional PH feature vectors on combined cancer cell and leukocyte positions. Representative images of the ROIs highlighted in the PCA plot are shown. Cancer cells are purple, leukocytes are mustard yellow. Images are 878µm x 878µm.

**Supplementary Figure 7 – linked to Figure 5.**
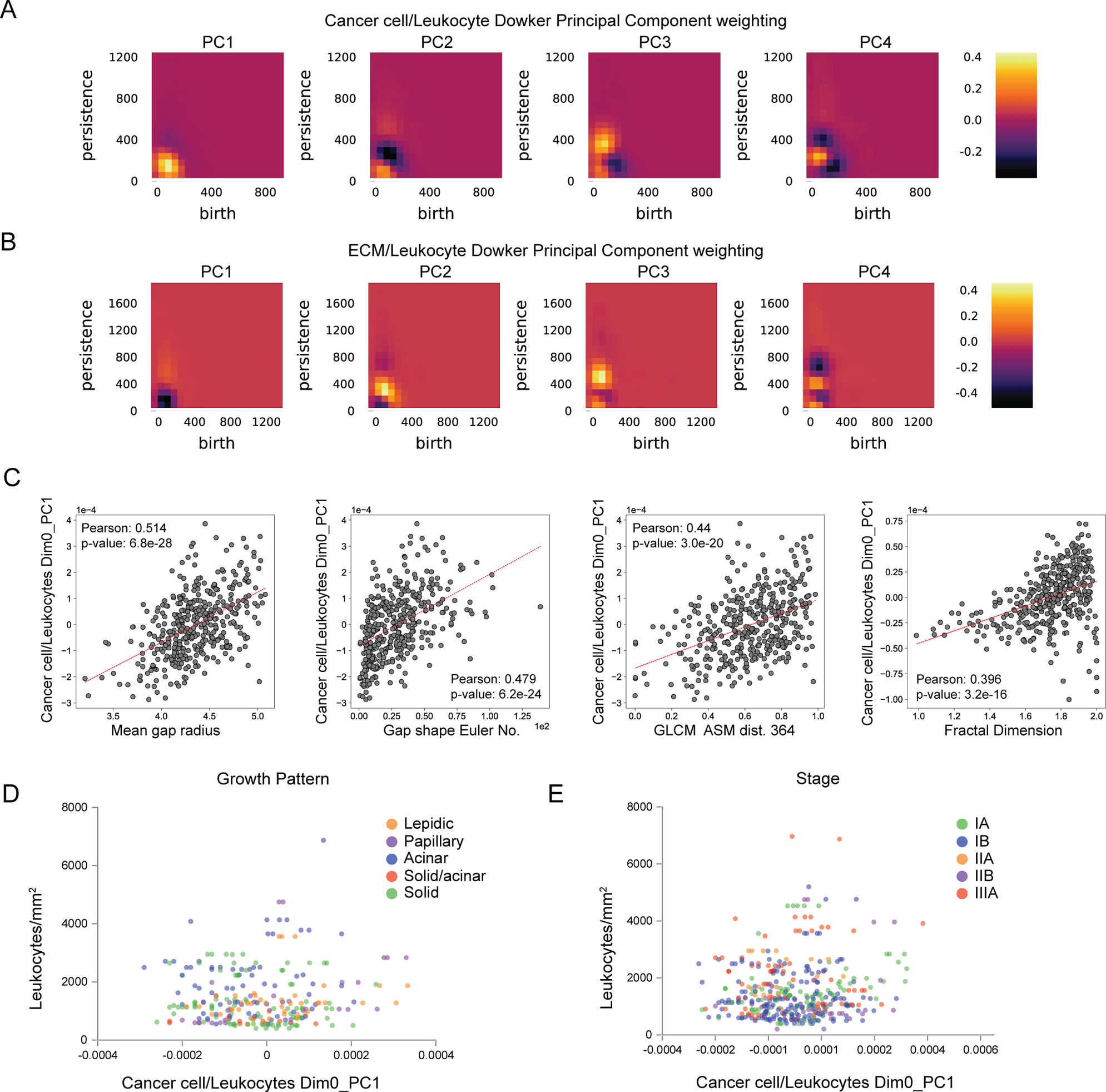
Dowker PH features and correlation of metrics. **A.** Visualization of the first four eigenvectors from PCA of 0-dimensional Dowker PH features of cancer cells and leukocytes (Fig. 5B) as persistence images. The dark regions indicate a lack of points in the persistence diagram in comparison to the mean, and the bright regions indicate an abundance of points in the persistence diagram in comparison to the mean. The first eigenvector indicates that ROIs whose persistence images with a high value in the yellow region will have a high PC1 coordinate. **B.** Visualization of the first four eigenvectors from PCA of 0-dimensional Dowker PH features of ECM and leukocytes (Fig. 5E) as persistence images. **C**. PC1 of 0-dimensional Dowker PH features of cancer cells and leukocytes (x-axis), leukocyte density (y-axis), and histological growth pattern (dot colours). **D**. Plot shows PC1 of 0-dimensional Dowker PH features of cancer cells and leukocytes (x-axis), leukocyte density (y-axis), and tumour stage (dot colours).

**Supplementary Figure 8 – linked to Figure 5.**
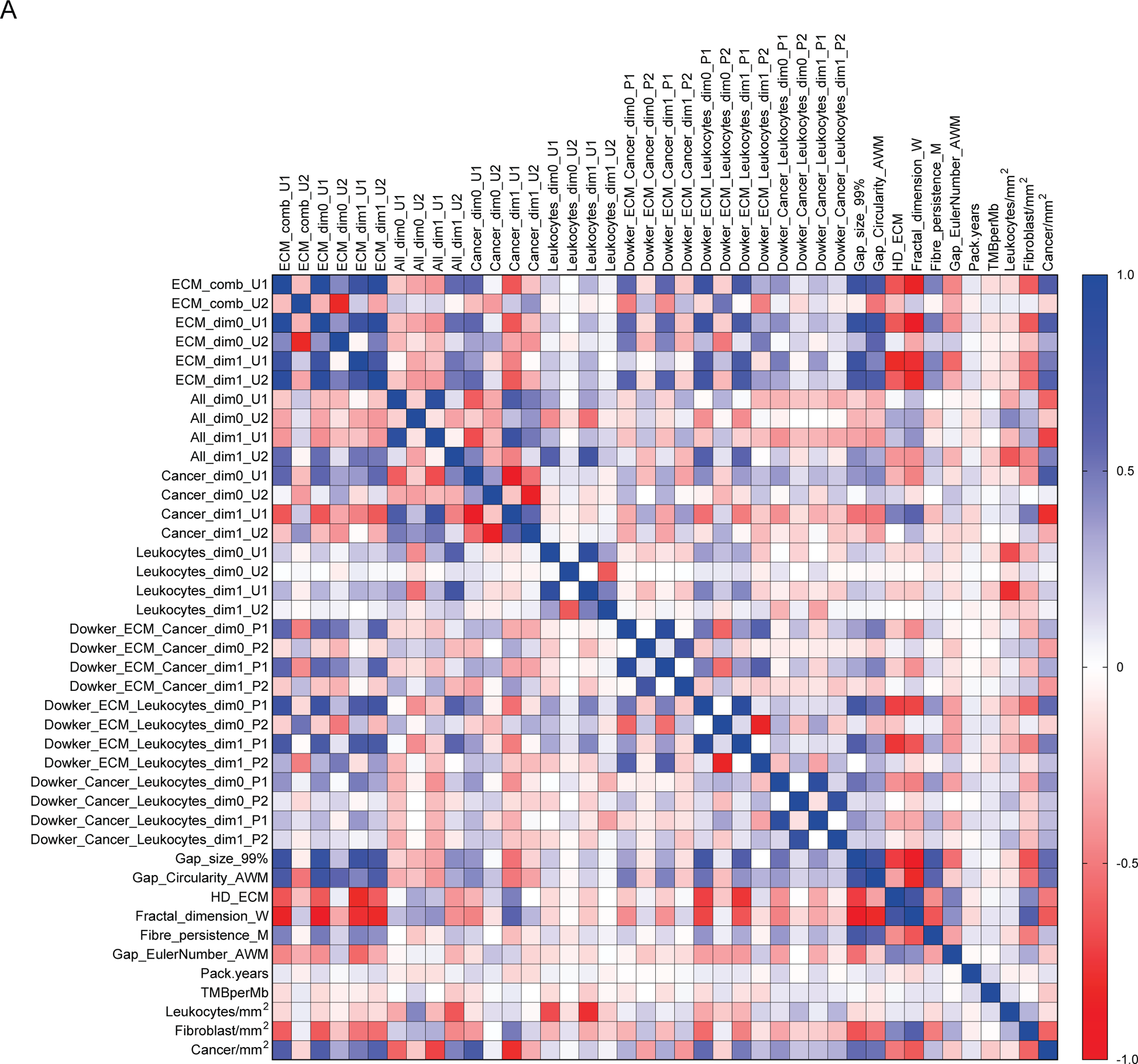
Plot shows the Pearson correlation of the indicated metrics. Blue indicates positive correlation, red indicates negative correlation.

**Supplementary Figure 9 – linked to Figure 7.**
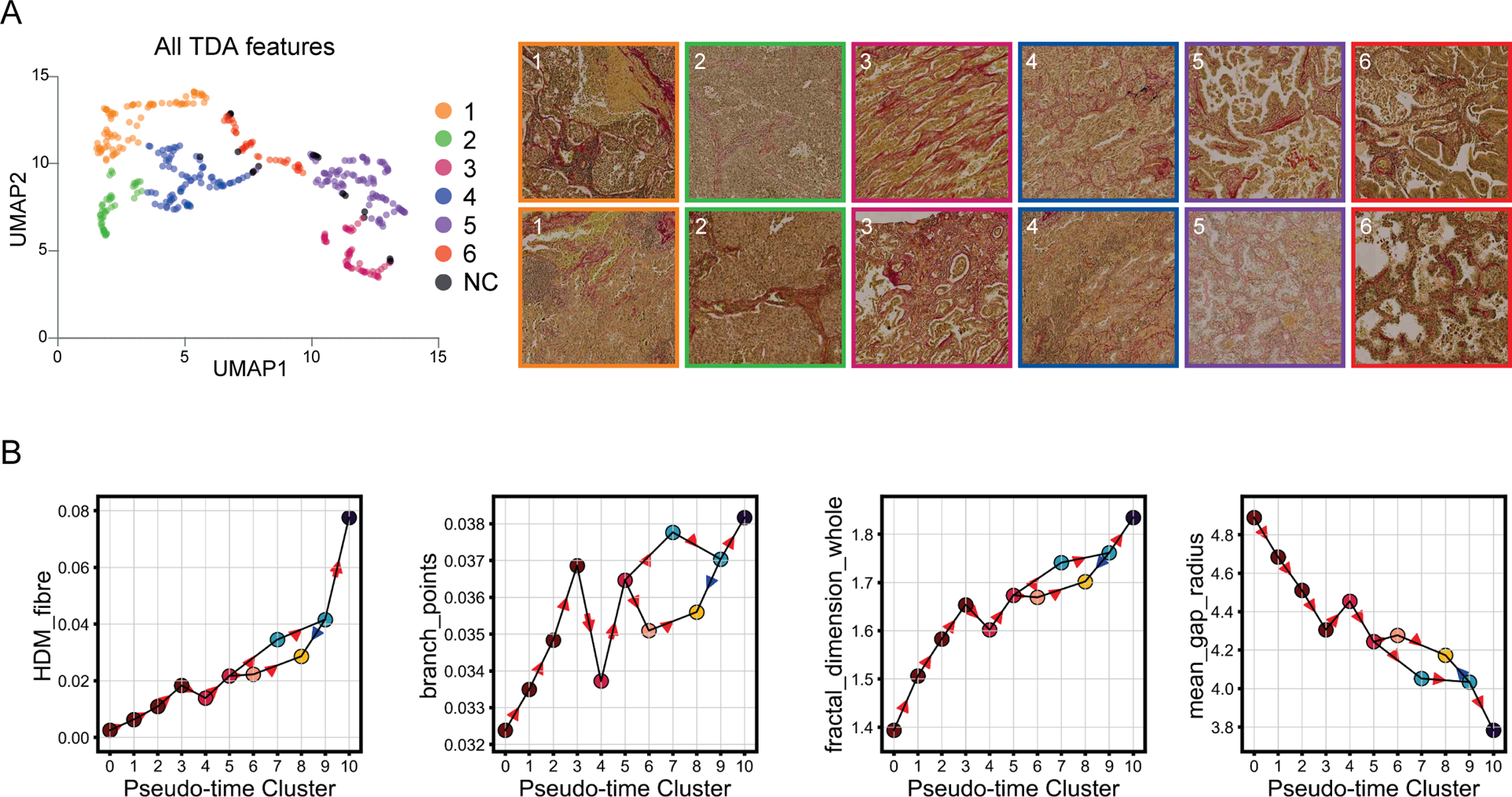
Integrated analysis of all features. **A**. (Left) UMAP generated on a concatenation of all PH features and Dowker PH features. (Right) Representative images from the six main clusters. **B**. Plots show how features vary as a function of pseudo-time cluster (x-axis).

## FUNDING ACKNOWLEDGEMENTS

The authors express their sincere thanks to The Mark Foundation for Cancer Research who have generously supported this work, and funded the contributions of I.H.R.Y. in full, and of E.C., D.N. and H.M.B. in part (MFCR ASPIRE 2022-0384).

E.S. and R.J. receive funding from the European Research Council (ERC Advanced Grant CAN_ORGANISE, Grant agreement number 101019366).

E.S. and A.R. are supported by the Francis Crick Institute which receives its core funding from Cancer Research UK (FC001144, FC001003), the UK Medical Research Council (FC001144, FC001003), and the Wellcome Trust (FC001144, FC001003).

C.S. is a Royal Society Napier Research Professor (RSRP\R\210001). His work is supported by the Francis Crick Institute that receives its core funding from Cancer Research UK (CC2041), the UK Medical Research Council (CC2041), and the Wellcome Trust (CC2041). For the purpose of Open Access, the author has applied a CC BY public copyright licence to any Author Accepted Manuscript version arising from this submission. C.S. is funded by Cancer Research UK (TRACERx (C11496/A17786), PEACE (C416/A21999) and CRUK Cancer Immunotherapy Catalyst Network); Cancer Research UK Lung Cancer Centre of Excellence (C11496/A30025); the Rosetrees Trust, Butterfield and Stoneygate Trusts; NovoNordisk Foundation (ID16584); Royal Society Professorship Enhancement Award (RP/EA/180007 & RF\ERE\231118); National Institute for Health Research (NIHR) University College London Hospitals Biomedical Research Centre; the Cancer Research UK-University College London Centre; Experimental Cancer Medicine Centre; the Breast Cancer Research Foundation (US) (BCRF-22-157); Cancer Research UK Early Detection and Diagnosis Primer Award (Grant EDDPMA-Nov21/100034); and The Mark Foundation for Cancer Research Aspire Award (Grant 21-029-ASP). C.S. was also supported by a Stand Up To Cancer-LUNGevity-American Lung Association Lung Cancer Interception Dream Team Translational Research Grant (Grant Number: SU2C-AACR-DT23-17 to S.M. Dubinett and A.E. Spira). Stand Up To Cancer is a division of the Entertainment Industry Foundation. Research grants are administered by the American Association for Cancer Research, the Scientific Partner of SU2C. C.S. is in receipt of an ERC Advanced Grant (PROTEUS) from the European Research Council under the European Union’s Horizon 2020 research and innovation programme (grant agreement no. 835297).

